# Characterization of two novel proteins involved in mitochondrial DNA anchoring

**DOI:** 10.1101/2020.07.22.215871

**Authors:** Simona Amodeo, Anneliese Hoffmann, Albert Fradera-Sola, Irina Bregy, Hélène Baudouin, Beat Haenni, Benoît Zuber, Falk Butter, Torsten Ochsenreiter

## Abstract

*Trypanosoma brucei* is a single celled eukaryotic parasite in the group of the Excavates. *T. brucei* cells harbor a single mitochondrion with a singular mitochondrial genome, that consists of a unique network of thousands of interwoven circular DNA molecule copies and is termed the kinetoplast DNA (kDNA). To ensure proper inheritance of the kDNA to the daughter cells the genome is linked to the basal body, the master organizer of the cell cycle in trypanosomes. The structure connecting the basal body and kDNA is termed the tripartite attachment complex (TAC). Using a combination of proteomics and RNAi (depletomics) we test the current model of hierarchical TAC assembly and identify TbmtHMG44 and Tb927.11.16120 as novel candidates of a structure that connects the TAC to the kDNA. Both proteins localize in the region of the unilateral filaments between TAC102 and the kDNA and depletion of each leads to a strong kDNA loss phenotype. TbmtHMG44 and Tb927.11.16120 stably associate with extracted flagella, even after DNase treatment however they do require the kDNA for initial assembly. Furthermore we demonstrate that recombinant Tb927.11.16120 is a DNA binding protein and thus a promising candidate to link the TAC to the kDNA.

## Introduction

*Trypanosoma brucei* is a single celled parasite that belongs to the Euglenozoa within the eukaryotic supergroup of the Excavates (Adl et al., 2019). The distinguishing feature of this group is the organization of its single mitochondrial genome into a large structure called the kinetoplast (Shapiro and Englund, 1995; Vickerman, 1977). In *T. brucei* it consists of 25 maxicircles, that encode 18 protein genes and two ribosomal RNAs. Twelve of the 18 genes are cryptogenes and require post transcriptional editing involving a large enzyme machinery (Editosome) and small trans-acting guide RNAs (Aphasizheva et al., 2020; Blum et al., 1990; Hajduk and Ochsenreiter, 2010; Read et al., 2016; Stuart et al., 2005). The guide RNAs are encoded on the 400 different minicircle species in the network coding for 1300 gRNA genes (Cooper et al., 2019). Overall there are 5-10′000 minicircles that are catenated with the maxicircles forming the kinetoplast DNA (kDNA) (Shapiro and Englund, 1995; Vickerman, 1977). Currently, more than 30 proteins have been described to be involved in the replication and segregation of the kDNA, while far fewer are required for this process in other eukaryotic systems (Jensen and Englund, 2012; Povelones, 2014; Schneider and Ochsenreiter, 2018). The model for kDNA replication predicts the release of minicircles into the region between the kDNA disc and the inner mitochondrial membrane, named the kinetoflagellar zone (KFZ). Here replication is thought to initiate, potentially through binding of the Universal Minicircle Binding Sequence Protein (UMSBP) or p38 to the universal minicircle sequence (UMS), a 12 nucleotide conserved sequence in the minicircles (Liu et al., 2006; Milman et al., 2007; Ryan et al., 1988). The processing then moves to the antipodal sites of the kDNA where two different primases, several polymerases and helicases and half a dozen proteins without obvious enzymatic function are involved in the completion of the replication process (for details see: (Bruhn et al., 2011; Concepción-Acevedo et al., 2018; Jensen and Englund, 2012; Klingbeil, 2004; Liu et al., 2009; Povelones, 2014).

The kDNA anchoring and segregation machinery of *T. brucei*, named the tripartite attachment complex (TAC), was discovered in 2003 and described by transmission electron microscopy analysis (Ogbadoyi, 2003). The TAC consists of three regions spanning three compartments of the cell (i) the exclusion zone filaments (EZF) that range from the base of the flagellum to the outer mitochondrial membrane (ii) the differentiated mitochondrial outer and inner membranes (DM) and (iii) the unilateral filaments (ULF) that range from the inner mitochondrial membrane to the kDNA (Ogbadoyi, 2003). The assembly of the TAC occurs *de novo*, from the base of the flagellum towards the kDNA, in a hierarchical manner, such that kDNA proximal components depend on the proper assembly of the kDNA distal components (Hoffmann et al., 2018; Schneider and Ochsenreiter, 2018). Our current model of the TAC includes 13 protein components (Schneider and Ochsenreiter, 2018). Four of these are localized to the EZF ((p197, BBA4, Mab22 and TAC65; (Bonhivers et al., 2008; Gheiratmand et al., 2013b; Käser et al., 2016)). The DM harbor four proteins of the TAC in the outer mitochondrial membrane (TAC60, TAC42, TAC40 and pATOM36, (Käser et al., 2016; Käser et al., 2017; Schnarwiler et al., 2014)) while one component is associated with the inner mitochondrial membrane (p166, (Ochsenreiter and Hajduk, 2006; Zhao et al., 2008)). The only known protein components of the ULF are TAC102 (Trikin et al., 2016) and TAP110 (Amodeo et al., 2020). While TAP110 is proximal to the kDNA when compared with TAC102, it does not seem to be essential for kDNA maintenance as other TAC proteins are (Schneider and Ochsenreiter, 2018). There are also a number of additional components that despite having multiple localizations are very likely part of the TAC. This includes for example the E2 subunit of the α-ketoglutarate dehydrogenase and the tubulin-binding cofactor C protein and AEP1 in the inner mitochondrial membrane (André et al., 2013; Sykes and Hajduk, 2013); (Ochsenreiter and Hajduk, 2006; Zhao et al., 2008)(André et al., 2013; Sykes and Hajduk, 2013). Aside from the proteins involved in the TAC are also several proteins that likely play a role in organization of the kDNA network and its anchoring to the TAC. Most prominently the kDNA associated proteins KAP3, KAP4 and KAP6. KAP3 was described as a histone H1 like protein in *Crithidia fasciculata*, while KAP6 is a HMG box containing protein described in *T. brucei* (Wang et al., 2014; Xu et al., 1996). Bloodstream form trypanosomes have a metabolically much reduced mitochondrion and mutants can be selected that are able to live without a mitochondrial genome. Dean and colleagues showed that the molecular basis for this capability a single point mutation in the γ subunit of the ATP synthase (Dean et al., 2013). This cell line provides a powerful tool that allows us to decipher direct from indirect effects on the mitochondrial genome maintenance machinery.

In this study we aimed to identify proteins responsible for linking the kDNA to the TAC and tried to characterize the organization of this region. For this we used a combination of proteomics, biochemical methods, RNAi and immunofluorescence microscopy.

## Results

### Interaction partners of TAC102

In an attempt to identify novel interaction partners of TAC102 we expressed a YFP tagged version of TAC102 in PCF cells, isolated cytoskeletons, sonicated the sample and immunoprecipitated TAC102 using an anti-GFP antibody. We identified 1523 proteins and among the top 50 enriched proteins, 21 were predicted/annotated to be mitochondrion or axonem associated. This included six already characterized TAC components: TAC102, TAP110, p166, TAC60, TAC65, and TAC40 (Amodeo et al., 2020; Käser et al., 2016; Käser et al., 2017; Schnarwiler et al., 2014; Trikin et al., 2016; Zhao et al., 2008). Also among the top 50 enriched proteins we detected the two very basic, potentially essential proteins, Tb927.9.5020 and Tb927.11.16120 that were predicted to be kinetoplast and mitochondrially localized, respectively (Table S1). Based on these findings we decided to characterize Tb927.9.5020 and Tb927.11.16120 in more detail. Tb927.11.16120 is a basic (pI = 10.0), conserved, and 68 kDa protein with a predicted mitochondrial targeting sequence at the N-terminus (Claros and Vincens, 1996), and was identified as component of the mitochondrial importome (Peikert et al., 2017). Tb927.9.5020 is a basic (pI 10.2), conserved, and 44 kDa large protein. It has a putative HMG-box domain at the N-terminus and thus we will refer to Tb927.9.5020 as TbmtHMG44. Homologs of both proteins can be found in the majority of the currently sequenced Kinetoplastea genomes but not in the Perkinsela sp. genome, an endosymbiotic kinetoplastid that lacks a basal body and flagellum (Tanifuji et al., 2017). Both proteins share a common evolutionary history (Figure S1) however, while we can identify a homologue of Tb927.11.16120 in the genome of the *Bodo saltans* (CUG90103.1) no homologue of TbmtHMG44 seems to be present in the free living kinetoplastid.

### TAC depletomics reveals organization of the kDNA-TAC interface

We used the current model of the hierarchical TAC organization to devise a proteomics strategy combining RNAi of a TAC protein or TAC associated component with the isolation of DNase treated flagella followed by quantitative mass spectrometry. From previous experiments we know that TAC proteins remain associated at the kDNA proximal end of the isolated flagella (Schneider and Ochsenreiter, 2018). Thus we expected that the closer the RNAi target is to the kDNA the fewer other proteins will be affected by the depletion of the target (see Figure 1A).

**Figure 1:**
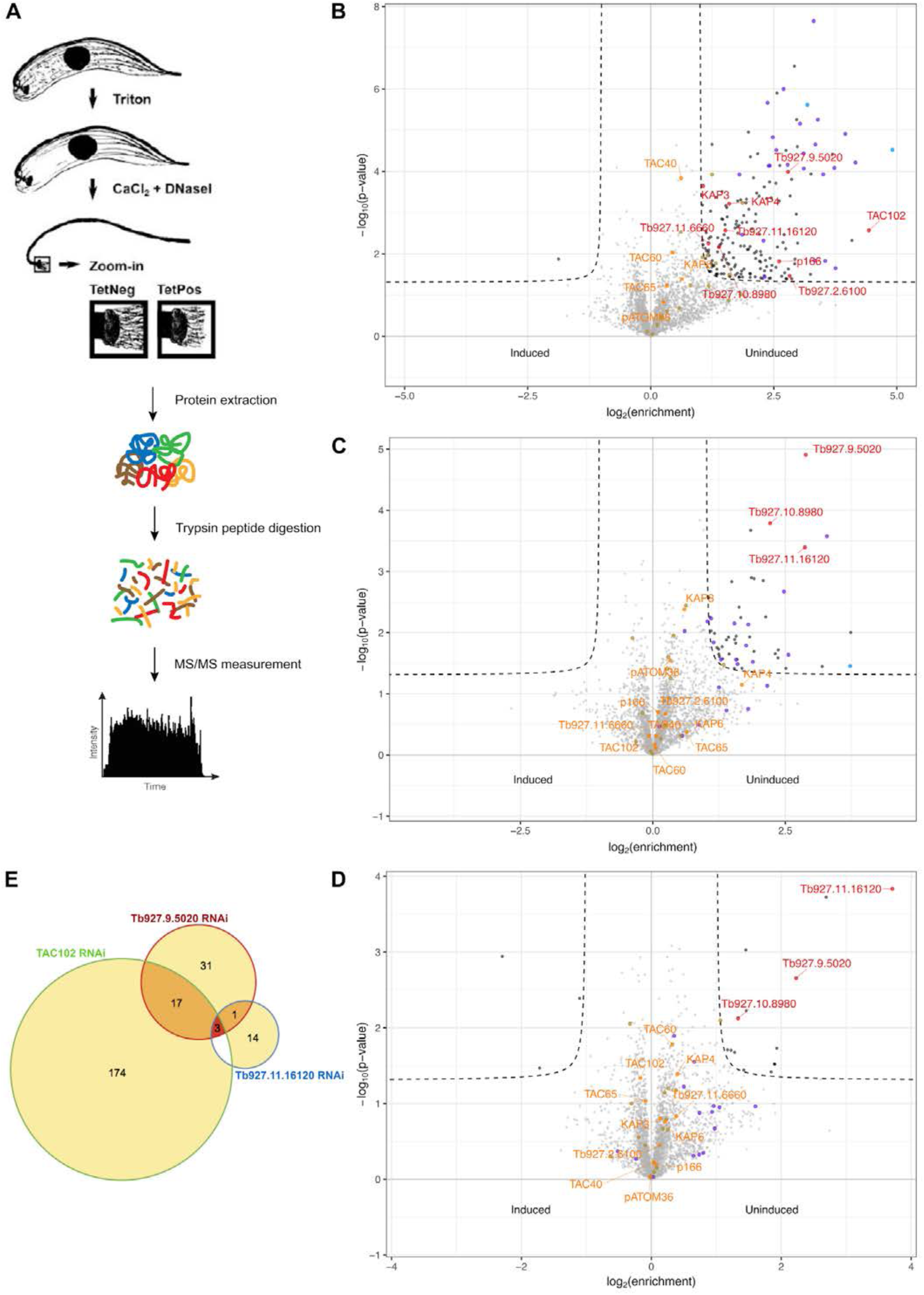
Mass spectrometry analysis of isolated flagella from BSF TAC102, Tb927.9.5020 and Tb927.11.16120 RNAi cells. **A)** Model of the experimental procedure of the flagellar extraction and mass spectrometry analysis. **B)** Volcano plot TAC102 RNAi. Identified proteins in uninduced flagella against the proteins in induced flagella at day three of the RNAi. The threshold was set as follows: p-value <0.05 and log2FoldChange >1 or <-1. **C)** Volcano plot Tb927.9.5020 RNAi. Same conditions as described in A). **D)** Volcano plot Tb927.11.16120 RNAi. Same conditions as described in A). **E)** Venn diagram of the proteins enriched in uninduced flagella from TAC102 RNAi, Tb927.9.5020 RNAi and Tb927.11.16120 RNAi. Highlighted in **blue**: oxidative phosphorylation factors. Highlighted in **purple**: mitochondrial gene expression factors. Highlighted in **khaki**: kDNA replication factors.

This approach allowed us to further test our current model of the hierarchical organization (see (Hoffmann et al., 2018; Schneider and Ochsenreiter, 2018)) (Figure 1A-E). We selected three proteins, TAC102, TbmtHMG44, and Tb927.11.16120. BSF cells expressing the RNAi constructs were used in flagellar extractions with DNase treatment followed by mass spectrometry. DNase treatment was required for a reliable sample preparation. The proteins that we identified in four replicates for the TAC102, the TbmtHMG44, and the Tb927.11.16120 RNAi experiment are shown in volcano plots (thresholds: p-value <0.05 and a fold change −1< log_2_ >1) (Figure 1B-D). Overall, the number of proteins identified by mass spectrometry was similar in all three experiments. However, while 194 proteins depend on the presence of TAC102, this number is lower for TbHMG44 (51, Figure S2, S3) and lowest for Tb927.11.16120, where only 15 interactors seem to depend on the presence of this protein (Figure S4). The only proteins that displayed dependencies in all three experiments were TbmtHMG44, Tb927.10.8980 and Tb927.11.16120 (Figure 1B-E). Out of the 194 proteins that were affected by TAC102 depletion >75% are predicted to be mitochondrially localized (Figure S5A). In the corresponding experiments targeting TbmtHMG44 and Tb927.11.16120 the fraction of proteins with predicted mitochondrial localization is around 60% and 30%, respectively (Figure S5B-C). Neither the depletion of TAC102 nor of TbmtHMG44 or Tb927.11.16120 changed the abundance of most characterized TAC components found in the flagellar extract (Figure 1B-D). One exception was the inner mitochondrial membrane protein p166, that was found to be decreased in abundance upon TAC102 RNAi (Figure 1B). This confirmed the previously observed interdependency of TAC102 and p166 (Hoffmann et al., 2018). Furthermore, the abundance of four kDNA associated proteins Tb927.11.6660, TbmtHMG44, Tb927.10.8980 and Tb927.11.16120 was affected by TAC102 depletion (Figure 1B). Tb927.2.6100 was previously described as a kDNA associated factor important for kDNA maintenance (Beck et al., 2013) while Tb927.11.6660 was recently identified as an interactor of TAC102 (Amodeo et al., 2020). Mass spectrometry also revealed the presence of 25 mitochondrial DNA replication factors in the DNase treated flagellar extracts (Table S2). They cover all stages of the replication process including minicircle release (TbTOPO2), replication initiation (UMSBP1/2) synthesis PolID, Primase 1 and Primase 2, as well as reattachment and gap closure (Pol beta, Pol beta PAK, MiRF172). The Primase 2 was decreased in abundance in all three depletomics experiments (Table S2), while Pol beta PAK, POLID, Primase 1, the kDNA associated proteins KAP3 and KAP4, the hypothetical protein Tb927.2.6100, the helicase TbPIF5 and the protease HslU2, were affected only in the TAC102 knockdown. Thus 33% of the detected replication factors seem to depend on the presence of TAC102, while only the Primase 2, required for minicircle replication initiation, depends on all three proteins (TAC102, Tb927.9.5020, Tb927.11.16120, Table S2).

### Tb927.11.16120 and TbmtHMG44 localize in the unilateral filament region of the TAC and are closer to the kDNA than TAC102

To localize Tb927.11.16120, we used an *in situ* PTP epitope tag in bloodstream form (BSF) and procyclic (PCF) *T. brucei* cells. By immunofluorescence microscopy Tb927.11.16120-PTP localized at the kDNA throughout the cell cycle in BSF as well as PCF cells (Figure S6A-C). Segregation of Tb927.11.16120 during kDNA replication occurred after separation of the basal bodies (Figure S6B, C arrowheads), and never prior to the division of the kDNA (Figure S6B, C arrowheads). Using STED super resolution microscopy we observed that Tb927.11.16120 and TAC102 signals partially overlapped, but that in most cases Tb927.11.16120 localized closer to the kDNA than TAC102 (Figure 2). Furthermore, we observed that during kDNA segregation the Tb927.11.16120 still formed a continuous structure at the time when TAC102 was already separated into two distinct signals (Figure 2, arrowheads). TbmtHMG44, was C-terminally HA-tagged *in situ* and localized by epifluorescence microscopy in close proximity to the kDNA in the region of the ULF (Figure 3A). Super-resolution microscopy showed that it localizes closer to the kDNA than TAC102 (Figure 3B). The signal for TbmtHMG44 was detectable alongside the entire face of the kDNA disk, but not overlapping with the signal of the kDNA or TAC102.

**Figure 2:**
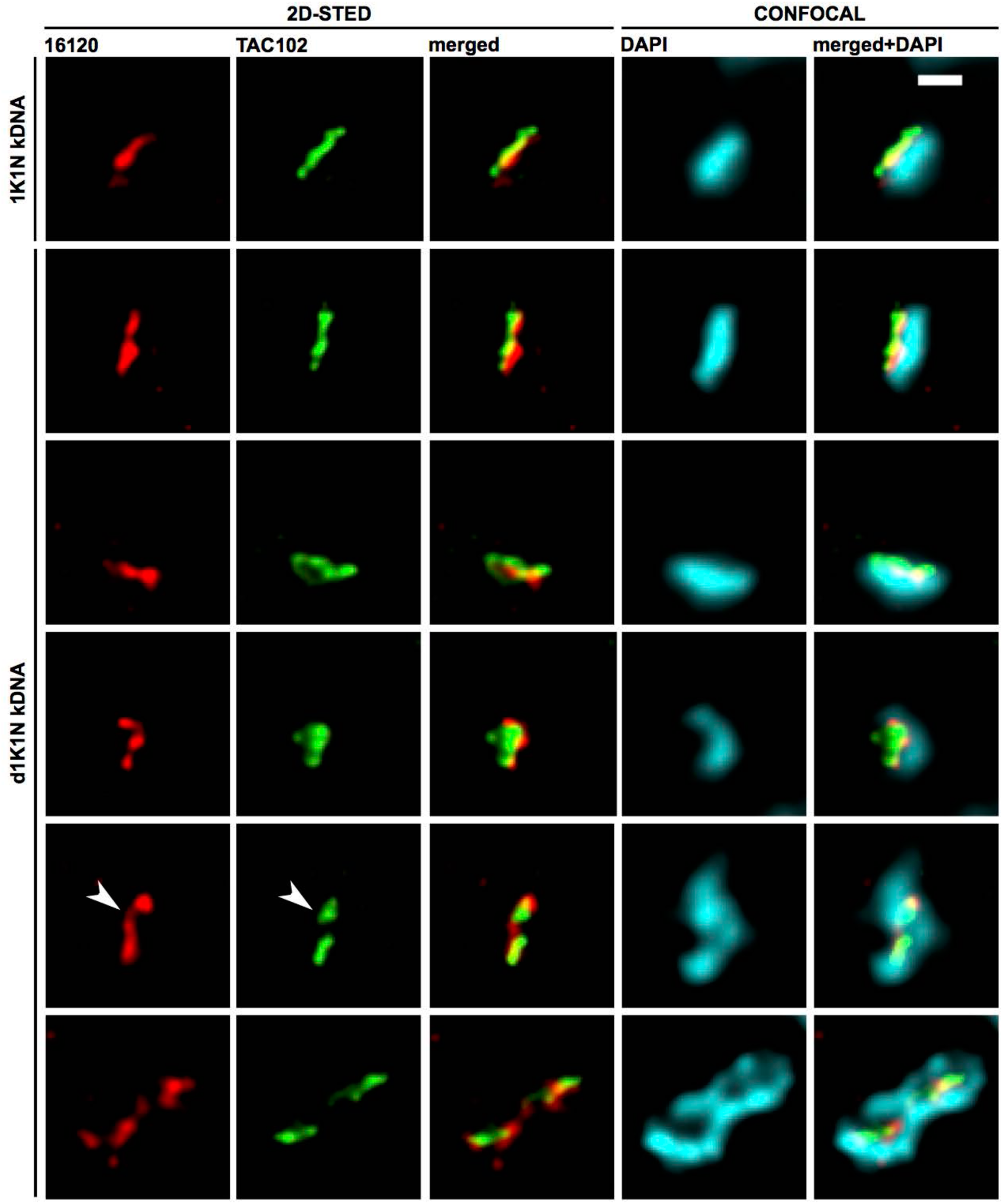
2D-STED immunofluorescence images of Tb927.11.16120-PTP (red) and TAC102 (green) at non-replicating and replicating kDNAs of BSF cells. The two proteins we detected with the same antibodies as described in the previous figures and acquired by 2D-STED. The DNA was stained with DAPI acquired with confocal microscopy. dK, replicating/duplicated kinetoplast; K, kinetoplast; N, nucleus. Scale bar 500 nm.

**Figure 3:**
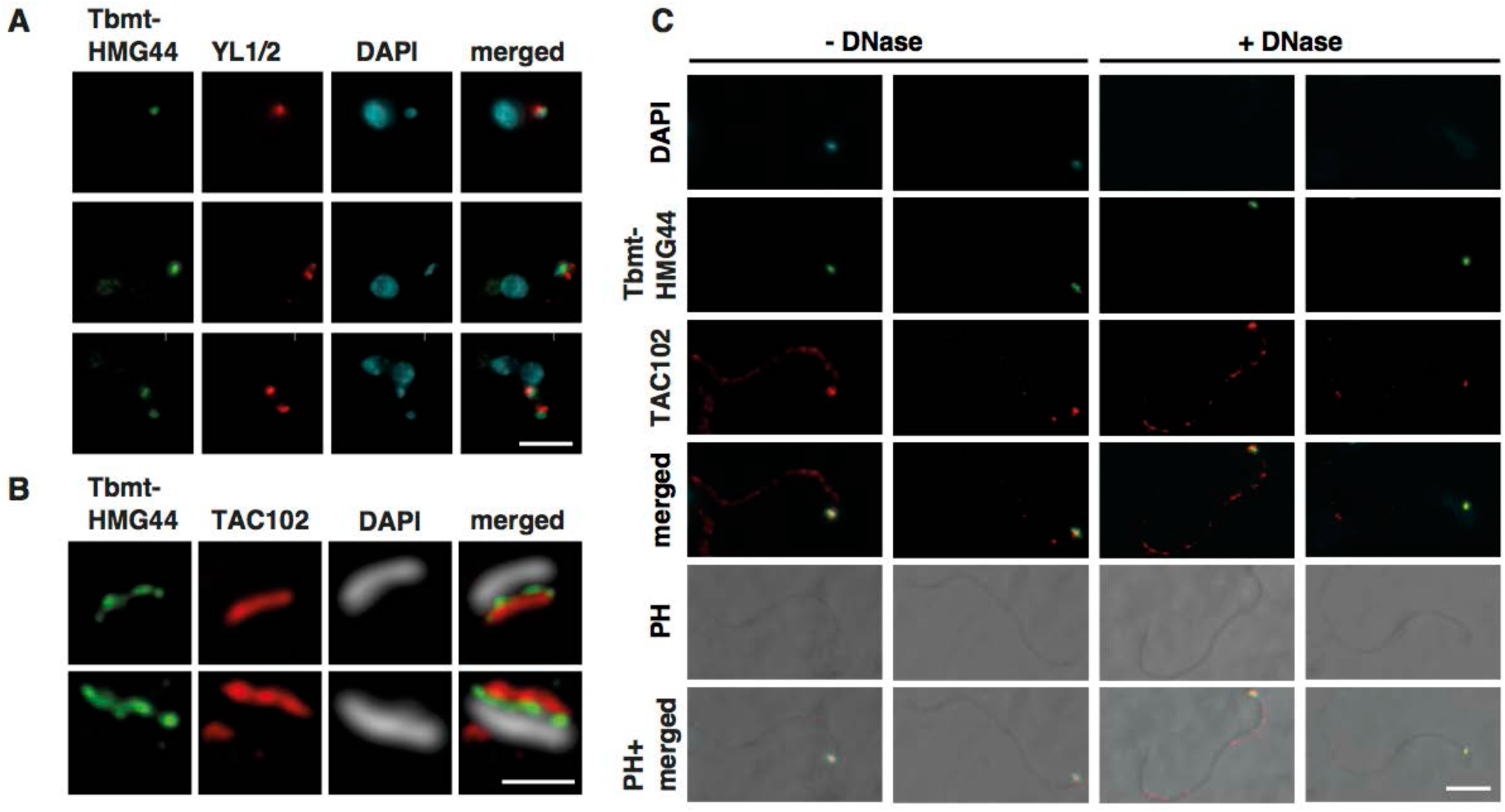
Localization of TbmtHMG44 in whole cells and isolated flagella. **A)** Representative images of HA-tagged TbmtHMG44 (green), YL1/2 (basal body, red) and DAPI (DNA, cyan) stained NYsm cells. Scale bar: 3 µm. **B)** Super-resolution images of TbmtHMG44 (green, HA) and TAC102 (red). The kDNA (white, DAPI) was acquired by confocal microscopy. Scale bar: 500 nm. **C)** DNase untreated (-DNase) and treated (+DNase) flagella extracts were stained with DAPI (DNA, cyan), TbmtHMG44 (HA, green) and TAC102 (red). Scale bar: 5 µm.

### TbmtHMG44 and Tb927.11.16120 remain stably associated with the TAC in DNase treated flagella but not in dyskinetoplastic cells

We recently showed that TAC102 as well as TAP110 remain at the flagellum even in dyskinetoplastic cells, supporting the model that TAC assembly is independent of kDNA (Amodeo et al., 2020; Hoffmann et al., 2018). To test whether this is also the case for TbmtHMG44 and Tb927.11.16120, we performed flagellar extractions in presence and absence of an endonulcease (DNase I). Like TAC102, both TbmtHMG44-HA and Tb927.11.16120-PTP remained attached to the flagellum, even after endonuclease treatment (Figure 3C, S6D). Furthermore, we described that the TAC is able to assemble in the absence of kDNA. The corresponding dyskinetoplastic cell line can be created through depletion of the TAC protein p197 (Gheiratmand et al., 2013a; Hoffmann et al., 2018) in the BSF γL262P cell line. Depletion of p197 disrupts the TAC, which leads to loss of the kDNA without death of the cell. After five days of p197 RNAi, all cells of the population are dyskinetoplastic and the cells can be released from RNAi (Hoffmann et al., 2018). In the course of recovery from p197 RNAi, the TAC reassembles and even replication factors like MiRF172 reassociate with the TAC in these cells (Amodeo et al., 2018). To test, whether the TAC alone is sufficient to localize TbmtHMG44 and Tb927.11.16120 to the region of the ULF we performed the p197 depletion experiment and monitored the localization of TbmtHMG44-HA (Figure 4) and Tb927.11.16120-PTP (Figure S7) by immunofluorescence microscopy. The basal bodies, which are not affected by p197 depletion, were detected by the YL1/2 antibody and served as marker for the region of the basal body relative to the TAC. As a TAC marker we used TAC102. At day five of p197 RNAi we verified that all cells had lost their kDNA (Figure 4A, S7C) and investigated the localization of TAC102, TbmtHMG44-HA, and Tb927.11.16120-PTP by epifluorescence microscopy (Figure 4B-D, S7B-E). A signal for TbmtHMG44-HA was detected in around 70% of uninduced cells. After five days of p197 depletion, about 80% of the cells had no detectable signal for TbmtHMG44-HA, while in about 20% of the cells TbmtHMG44-HA was much less abundant or at the wrong position in the mitochondrion (Figure 4B). In case of Tb927.11.16120-PTP, 100% of the cells showed a signal prior to p197 depletion. After five days of p197 RNAi, we observed that approximately 80% of the cells had no detectable signal for Tb927.11.16120-PTP, and in the other 20% of the cells we only observed a weak signal for Tb927.11.16120-PTP or it was at the wrong position in the mitochondrion (Figure S7E). In dyskinetoplastic cells, three days post recovery from p197 RNAi in almost 100% of the cells TAC102 was detected at the correct position (Figure 4B, S7D), however only 20% and 10% of the cells showed a wild type signal for TbmtHMG44 (Figure 4C) and Tb927.11.16120 (Figure S7E), respectively. Thus the TAC is not sufficient to correctly localize TbmtHMG44 and Tb927.11.16120 to the inner ULF. Both proteins require the kDNA for proper localization.

**Figure 4:**
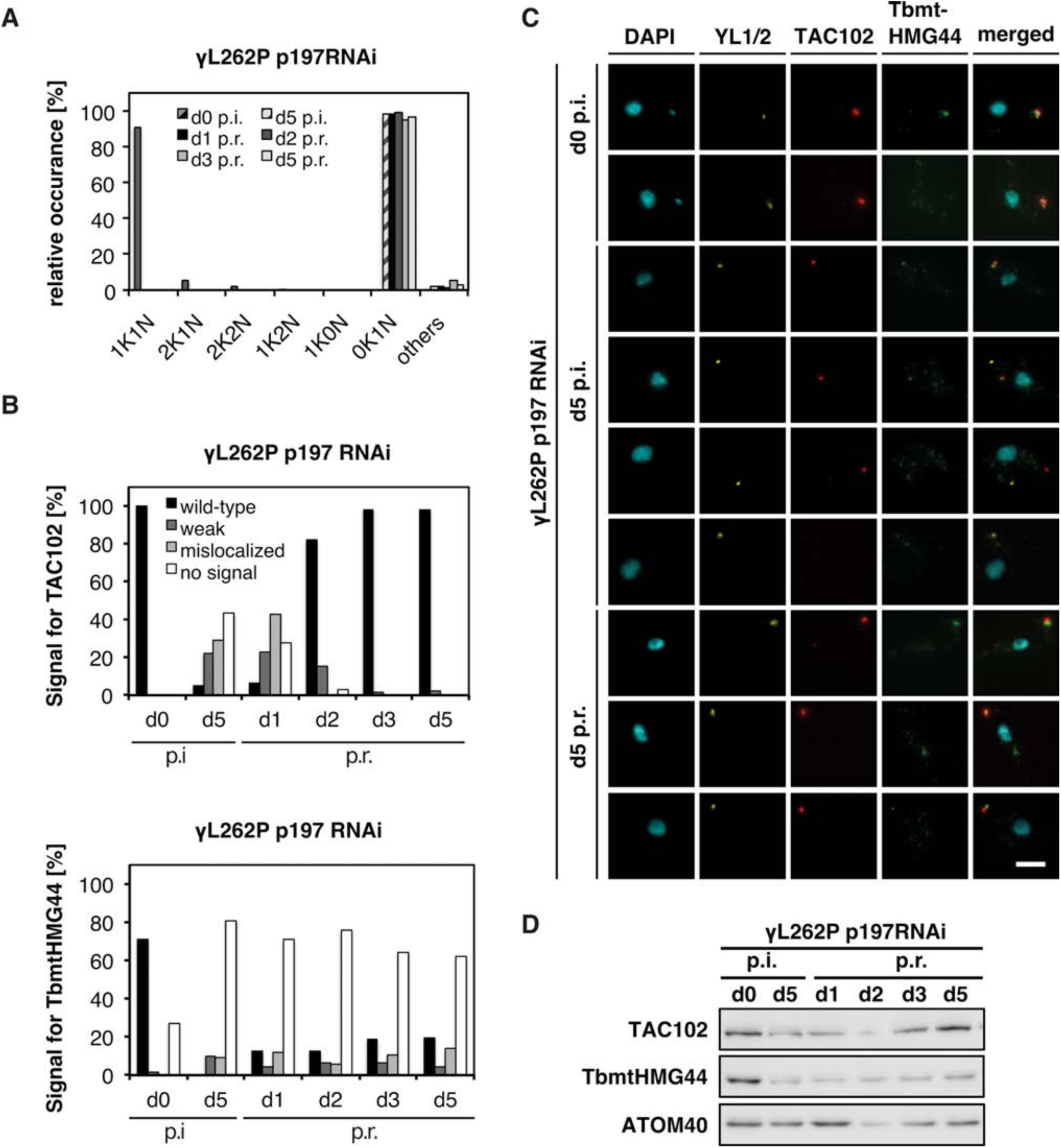
Recovery of TbmtHMG44-HA signals in γL262P p197 RNAi cells. **A)** Cell cycle stages of uninduced and 5 days induced (day 5 p.i.) cells as well as on day 1, 2, 3, 5 after removing tet (post recovery, p.r.) (N≥92). **B)** Quantitative analysis of TAC102 and TbmtHMG44HA in YL1/2 positive cells. The counting was performed in the same cells. A black bar indicates a wild-type localization, a grey bar depicts a weak signal, a light grey bar indicates a mislocalized signal, and a white bar no signal (N≥92). **C)** Representative images of the quantification in B). Cytoskeleton extracted cells were stained with DAPI (DNA, cyan), YL1/2 (basal body, yellow), TbmtHMG44 (HA, green) and TAC102 (red) after 0 and 5 days of tet induction and 5 days post recovery **D)** Protein abundance of TAC102 and TbmtHMG44-HA in uninduced, induced (day 5 p.i.) and cells after re-expression of p197 (p.r., post recovery). ATOM40 was used as a loading control. Scale bar 3 µm.

### Depletion of Tb927.11.16120 or TbmtHMG44 causes strong kDNA loss

To study the function of Tb927.11.16120 and TbmtHMG44, we performed RNAi targeting the open reading frames in BSF cells and monitored growth, kDNA phenotype and the effect on TAC assembly. After Tb927.11.16120 RNAi induction, cells grew at normal rates until day four, when a growth defect became visible (Figure 5A). The population started to collapse at day five post induction of Tb927.11.16120 RNAi. A northern blot revealed that the Tb927.11.16120 mRNA was decreased by 70% at day three post induction (Figure 5A, inset). We then also analyzed the impact of Tb927.11.16120 depletion on the kDNA, and we detected an accumulation of cells with small kDNA (s1K1N) and no kDNA (0K1N) (Figure 5B). At day three post induction around 60% of the cells had no kDNA and around 20% of the cells had small kDNAs (Figure 5B, C). We also observed few missegregated kDNAs (Figure 5B), a typical characteristic for TAC components (Schneider and Ochsenreiter, 2018). RNAi targeting TbmtHMG44 led to a very similar phenotype. The cells grew slower at day four and the population immediately collapsed afterwards (Figure S8A). qPCR revealed a decrease of TbmtHMG44 mRNA by 70% at day three of the RNAi (Figure S8A, inset). Furthermore, at day three post TbmtHMG44 RNAi induction around 85% of the cells had lost their kDNA (Figure S8B, C). To test whether the depletion of TbmtHMG44 impacts mitochondrial morphology, we also stained the cells for the mitochondrial HSP70 protein. No obvious morphological changes of the mitochondrion were detected three days post TbmtHMG44 RNAi induction (Figure S8D).

**Figure 5.**
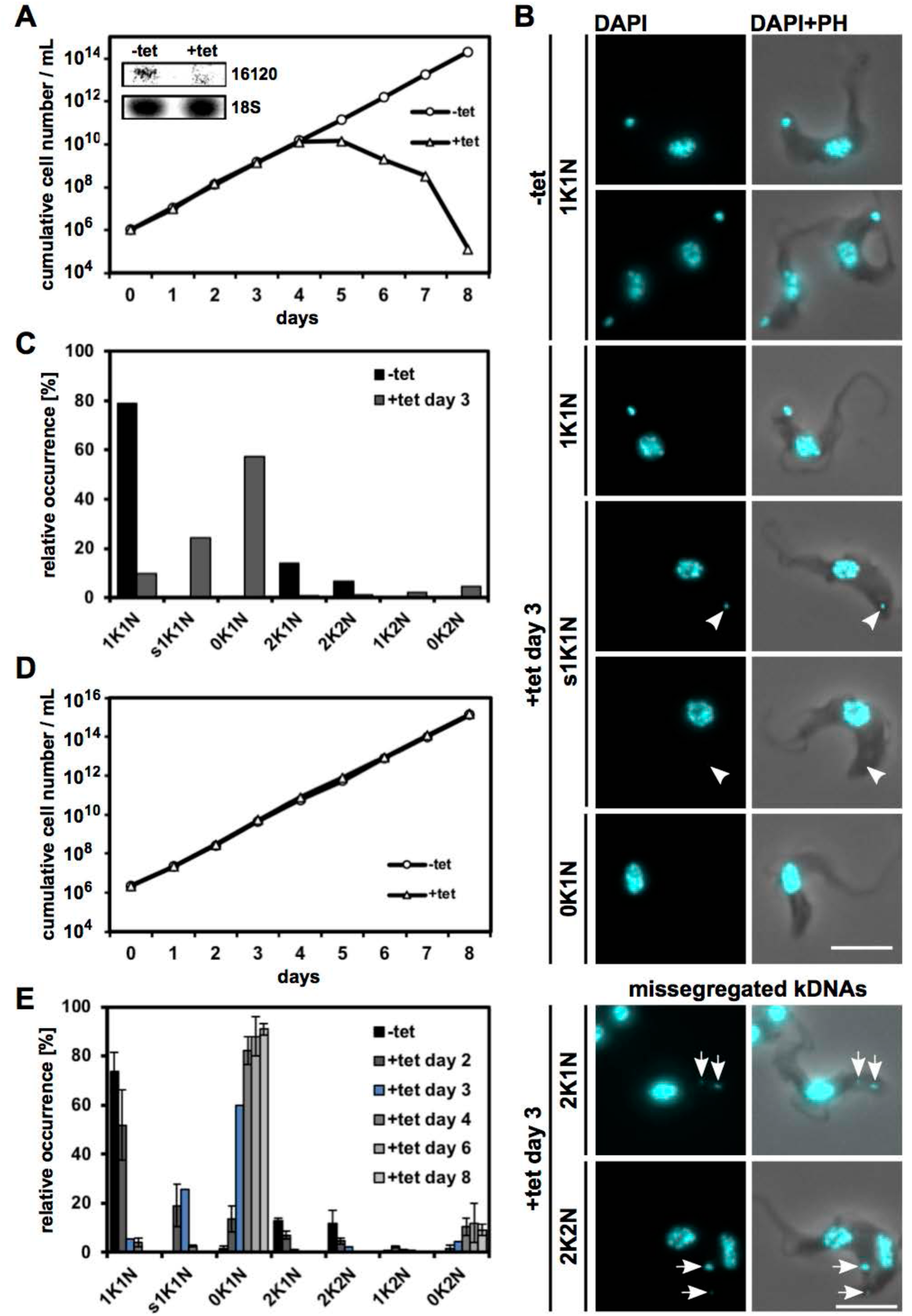
Phenotype upon depletion of Tb927.11.16120 in BSF cells. **A)** Growth curve of Tb927.11.16120 RNAi expressing cells. The inset depicts a northern blot, showing knockdown of Tb927.11.16120 mRNA at day three post induction. 18S rRNA serves as a loading control. **B)** Representative fluorescence microscopy images of uninduced cells and cells at day three post induction. The nucleus and the kDNA were stained with DAPI. Arrowheads point to small kDNA, arrows to mis-segregated kDNAs. **C)** Quantification of the relative occurrence of kDNA discs and nuclei in Tb927.11.16120 RNAi induced (+tet day 3) and uninduced cells (−tet) (n≥100 for each condition). **D)** Growth curve of Tb927.11.16120 RNAi expressing γL262P BSF cells. **E)** Quantification of the relative occurrence of kDNA discs and nuclei in Tb927.11.16120 RNAi expressing γL262P BSF cells (n≥100 for each condition). K, kDNA; N, nucleus; PH, phase contrast; sK, small kDNA. Scale bar: 5 μm.

We then targeted Tb927.11.16120 and TbmtHMG44 by RNAi in a cell line which is able to survive the loss of kDNA due a mutation in the γ subunit of the ATP synthase (γL262P, (Dean et al., 2013)). The growth curve was performed during the course of eight days and we did not observe any effect on growth and viability of the cells during Tb927.11.16120 RNAi (Figure 5D) nor TbmtHMG44 RNAi (Figure S8F). We also quantified the kDNA loss and small kDNAs and observed the same effect on the kDNA as described above (Figure 5E, S8E, G). Together this indicates that neither Tb927.11.16120, nor TbmtHMG44, are essential for γL262P cells and therefore both proteins are likely to exclusively function in kDNA maintenance.

To verify that the tagged version of TbmtHMG44 is indeed functional we performed RNAi against the 3’UTR of the enodgenous TbmtHMG44 in the background of a cell line expressing TbmtHMG44-HA. After three days of TbmtHMG44 depletion we did not detect any loss of kDNA or change in localization of the tagged TbmtHMG44, however there was a weak growth defect visible (Figure S9A-C). This indicates that the HA-tagged version of TbmtHMG44 is mainly functional, as it rescues the kDNA loss phenotype.

### Depletion of TbmtHMG44 and Tb927.11.16120 affect kDNA network composition and structure

Since the depletion of TbmtHMG44 and Tb927.11.16120 led to kDNA loss, we wanted to further characterize the effect on the network structure, the different circle types (mini- and maxicircles) and minicircle replication intermediates (covalently closed (CC) circles prior to replication vs nicked and gapped (N/G) circles after completion of replication). To assess the effect on mini- and maxicircles, we used Southern blot analysis. Total mini- and maxicircle content can be analyzed when using HindIII and XbaI digested DNA extracted from the RNAi cell lines. We extracted DNA from the γL262P TbmtHMG44 RNAi and the γL262P Tb927.11.16120 RNAi cell lines, resolved them on a agarose gel, transferred the DNA to a membrane and probed for mini-and maxicircles. The amount of minicircles increased to about 150% at day two of Tb927.11.16120 depletion and the maxicircle content even increased to around 200% (Figure 6A, B). Depletion of TbmtHMG44 let to an increase of minicircles at day one only, while at day two, the minicircle content decrease to around 70% (Figure S10A, B). For the maxicircles a similar increase (as for Tb927.11.16120 RNAi) to around 230% (p <0.02) was observed at day two of the RNAi (Figure S10A, B). For Tb927.11.16120 depleted cells the mini- and maxicircle content was then massively decreased at day four post induction (Figure 6A, B). We did not assess the mini- and maxicircle content at day four of TbmtHMG44 RNAi but at day three observed a massive decrease of minicircles already, while there was still an increased content of maxicircles detectable (around 180%) (Figure S10A, B).

**Figure 6.**
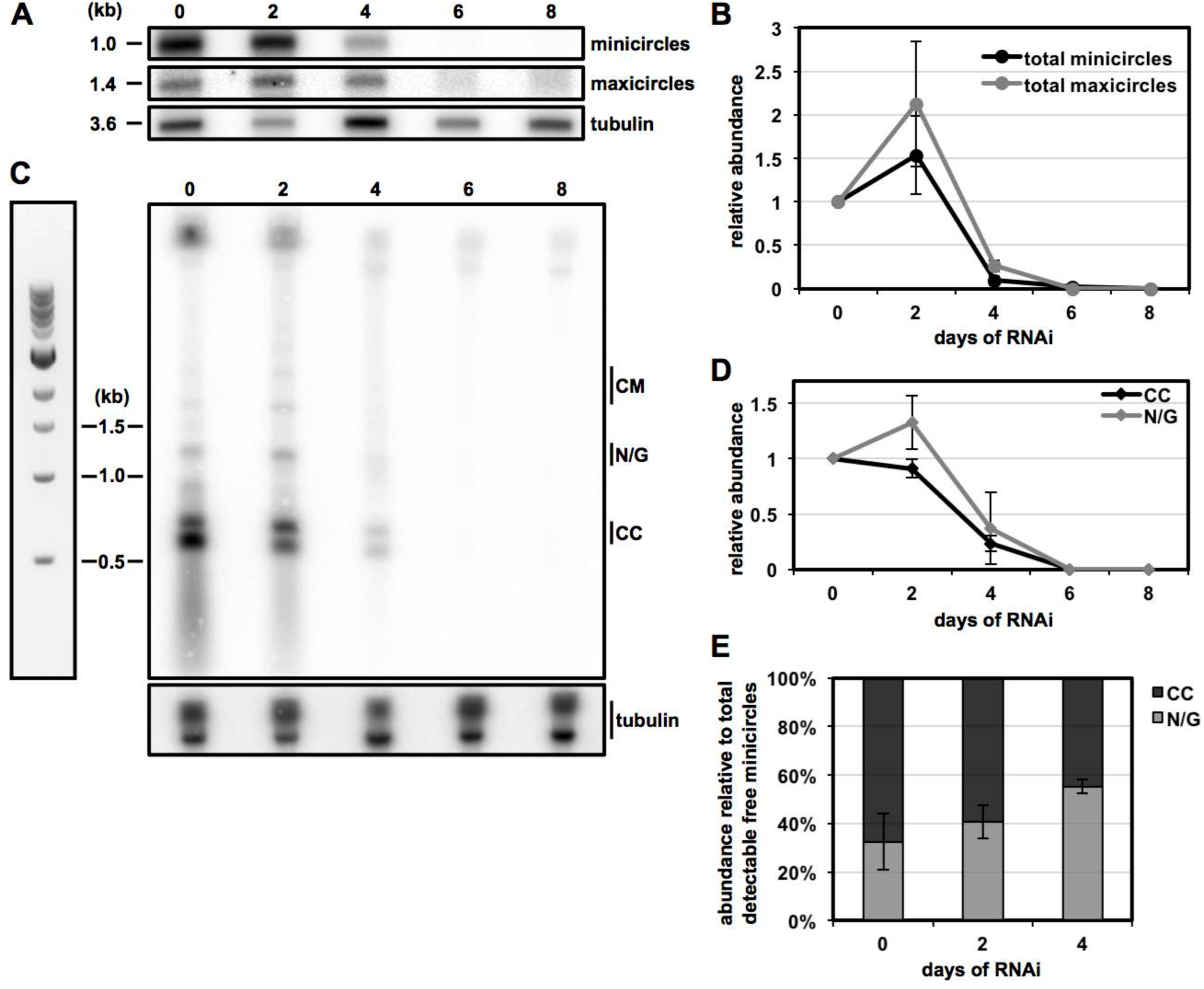
Effect of Tb927.11.16120 depletion on the abundance of kDNA and kDNA replication intermediates. **A)** Detection of total mini- and maxicircle by Southern blot. Total DNA (isolated from Tb927.11.16120 RNAi expressing γL262P BSF cells), digested with HindIII and XbaI, from either uninduced (0) or cells from different days post induction (day 2, 4, 6, 8) was used. Linearized minicircles, a 1.4kb maxicircle fragment and a 3.6kb fragment of the tubulin inter-genic region were detected with radioactive probes. **B)** Quantification of minicircle and maxicircle abundance as exemplified in A) (n=3). **C)** Detection of free minicircle replication intermediates by Southern blot as described in A) but without treatment of total DNA with restriction enzymes. **D)** Quantification of covalently closed and nicked/gapped free minicircles abundance as exemplified in C) (n=3). **E)** Quantification of the relative portions of covalently closed and nicked/gapped minicircles in the total of free minicircles (here just covalently closed+nicked/gapped) (n=2).

Free minicircle replication intermediates released for replication (CC and N/G) can be analyzed by Southern blot when using undigested DNA extracted from the RNAi cell lines. At day two of Tb927.11.16120 RNAi, the CC, not yet replicated, minicircles were detected with almost the same abundance as in non-induced cells, whereas at day four of the RNAi the CC level had dropped to about 20% (Figure 6C, D). The N/G, replicated but not yet reattached, minicircles in contrast increased to around 130% at day two of Tb927.11.16120 depletion and then the abundance rapidly decreased to around 40% at day four of the RNAi (Figure 6C, D). The amount of N/G minicircles relative to the total of N/G and CC minicircles, was increasing over the course of induction (Figure 6F). Knockdown of TbmtHMG44, in contrast, led to a constant loss of the CC minicircles (around 40% of CC were left at day three post induction) (Figure S10C, D). Similarly, depletion of TbmtHMG44 also led to a more or less constant loss of N/G minicircles (around 60% of N/G were lost at day three post induction) (Figure S10C, D). In addition, we also detected a slower migrating minicircles species that are probably catenated minicircles (CM) at day two of the TbmtHMG44 RNAi (Figure S10C).

To analyze the ultrastructure of the kDNA, we performed transmission electron microscopy (TEM) on ultrathin sections of embedded trypanosomes (Figure 7). When TbmtHMG44 was depleted we detected, besides cells without kDNA and small kDNAs, much weaker stained kDNAs without the typical striated appearance that seemed to have lost the compact disc-like structure (Figure 7).

**Figure 7:**
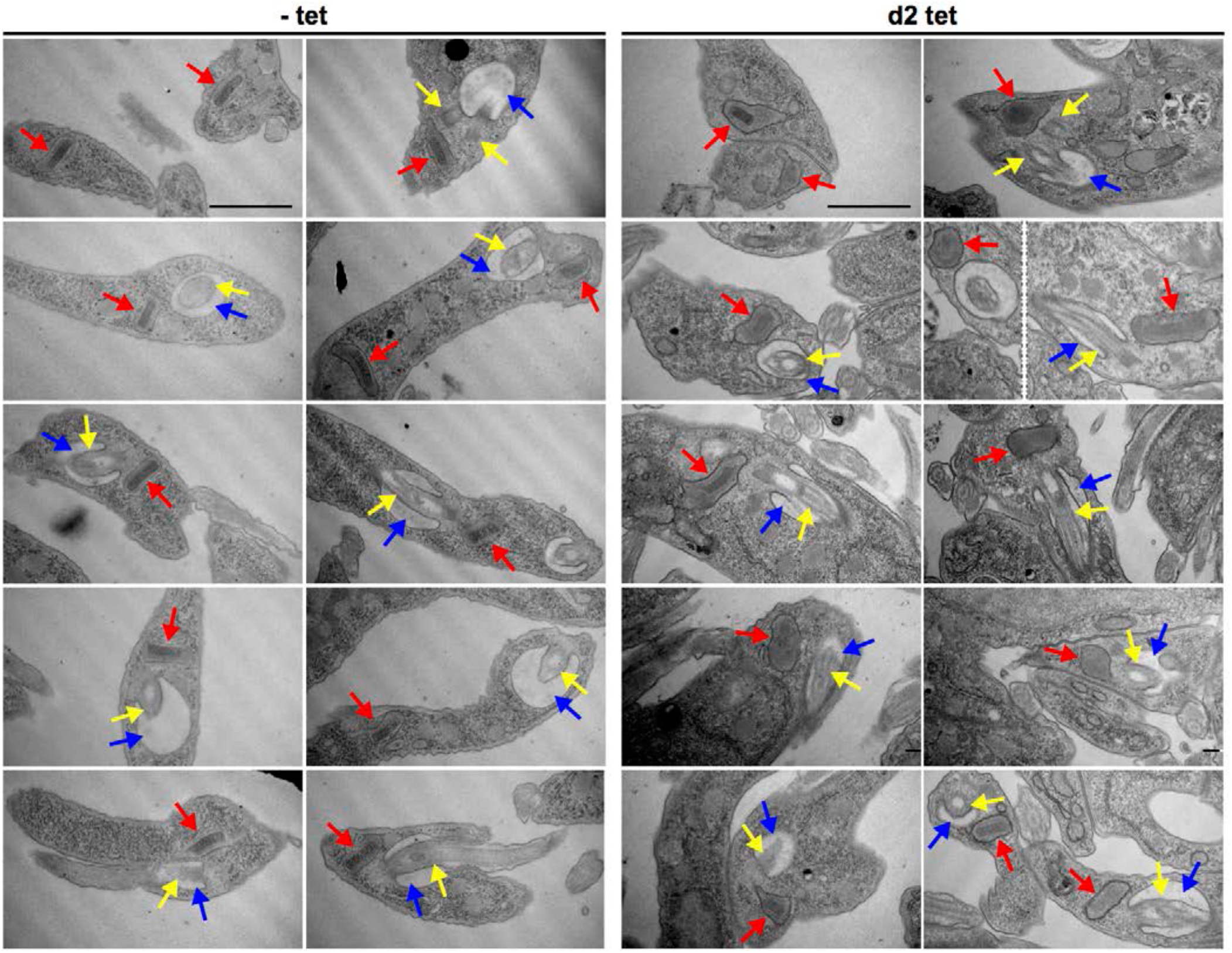
TEM analysis of TbmtHMG44 RNAi BSF cell. Representative images of the ultrastructure of the kDNA of TbmtHMG44 RNAi cells revealed by TEM. On the left side kDNAs from uninduced cells (- tet) are shown. Representative images of kDNAs from cells at day two post induction (d2 tet) are shown on the right side. Red arrows point to the kDNA in the kDNA pocket, the yellow arrows to the flagellum or base of flagellum and the blue arrows to the flagellar pocket. Scale bar = 1 μm.

### Depletion of TbmtHMG44 and Tb927.11.16120 do not lead to loss of TAC102

The 2D-STED microscopy analysis revealed that both, TbmtHMG44 and Tb927.11.16120 localize closer to the kDNA than TAC102. Based on the current model of TAC assembly (Hoffmann et al., 2018; Schneider and Ochsenreiter, 2018), we therefore would expect TAC102 to remain associated with the TAC even in the absence of TbmtHMG44 or Tb927.11.16120. The mass spectrometry analysis did already reveal that TAC102 remained associated with the TAC upon depletion of TbmtHMG44 as well as depletion of Tb927.11.16120 (Figure 1). We wanted to verify this by immunofluorescence microscopy. In general we did not observe a loss of TAC102 upon Tb927.11.16120 or TbmtHMG44 depletion (Figure 8A, B, S11A, B), however about 20% of the cells without kDNA showed a weaker signal of TAC102 when Tb927.11.16120 was depleted and about 5% of 0K1N cells had lost the signal for TAC102 (Figure S11B). Quantification of the kDNA loss phenotype showed results similar to our previous observations (Figure S11C) (Trikin et al., 2016). Similarly, TbmtHMG44 depletion did not have a major impact on TAC102 localization (Figure 8A), however in a few cells TAC102 signal intensity was decreased (Figure 8B, C).

**Figure 8:**
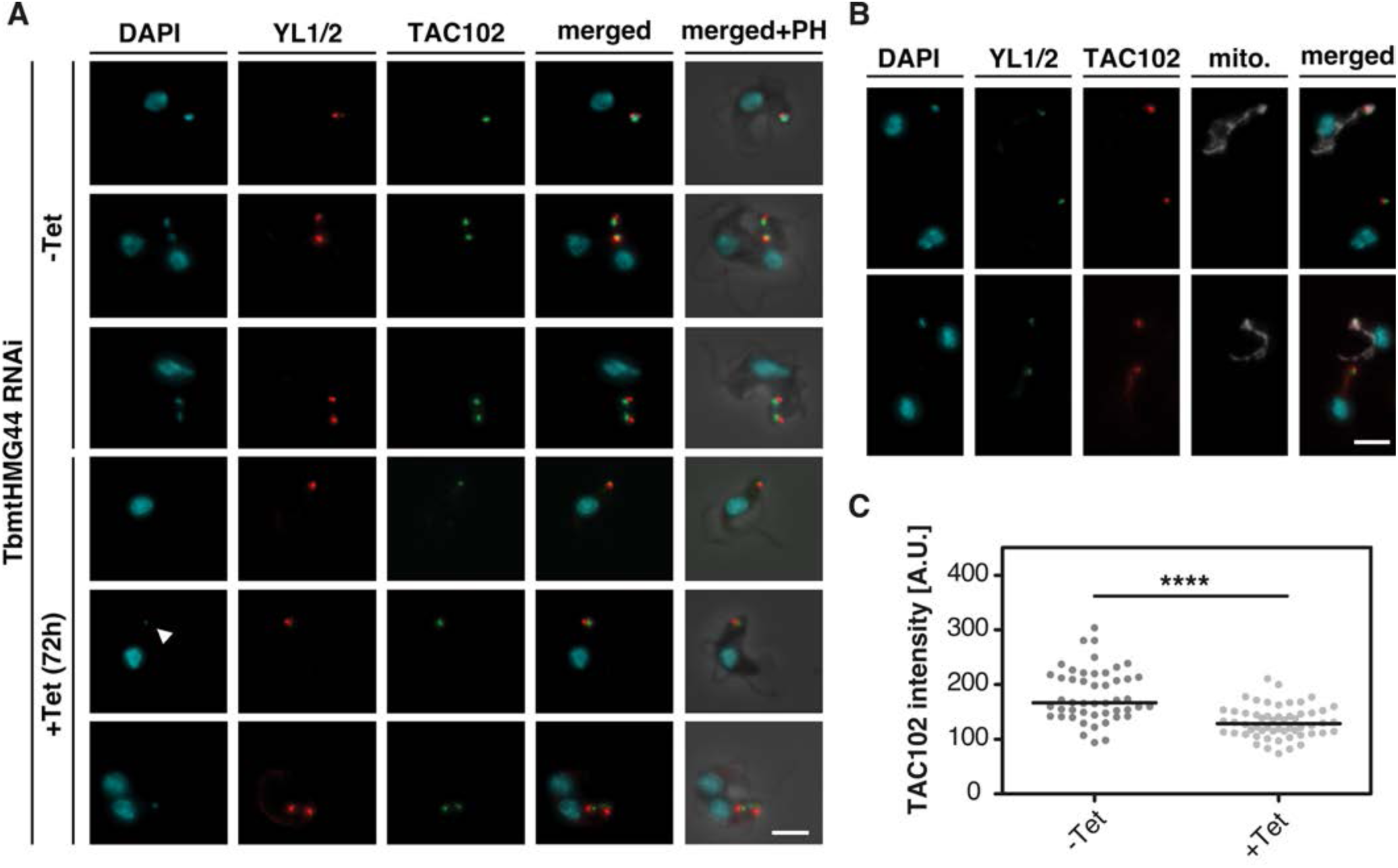
TAC102 signal upon TbmtHMG44 depletion in NYsm cells. **A)** Representative images of DAPI (cyan), YL1/2 (basal body, red) and α-TAC102 (green) stained cells either TbmtHMG44 uninduced (-tet) or 72 h induced (+tet). Merged and phase contrast (Ph) images are shown next to it. Arrowhead points towards a small kDNA. **B)** Representative images of mixed uninduced (MitoTracker (mitochondrion, white) stained) and induced cells stained with α-TAC102 (red) and YL1/2 (green, basal body). **C)** Quantitative TAC102 intensity measurements of the acquired pictures from B) (N_-tet_=49; N_+tet_=56). The p-value was calculated to perform significance measurements (two-tailed heteroscedastic t-test; **** p≤0.0001). Scale bar 3 µm

### Depletion of TAC102 leads to loss of Tb927.11.16120

The mass spectrometry analysis did reveal that TbmtHMG44 as well as Tb927.11.16120 were depleted from TAC102 RNAi induced, isolated flagella. We also wanted to verify this by immunofluorescence microscopy and therefore monitored whether TAC102 is required for localization of Tb927.11.16120. For this we used RNAi depletion of TAC102 and monitored the effect on Tb927.11.16120. We quantified immunofluorescence microscopy data of ≥100 TAC102 RNAi induced / non-induced cells (Figure 9A). As a control we assessed the kDNA loss phenotype upon TAC102 RNAi, which was comparable to the previously obtained data (Trikin et al., 2016) (Figure 9B). Two days post TAC102 RNAi induction 80% of the cells had lost the kDNA and showed no signal for TAC102 and Tb927.11.16120-PTP (Figure 9C, D). In cells with small kDNAs (s1K1N, Figure 9A, arrowhead) or big kDNAs (b1K1N, Figure 9A, arrow) a weak signal for TAC102 and Tb927.11.16120 was sometimes observed.

**Figure 9.**
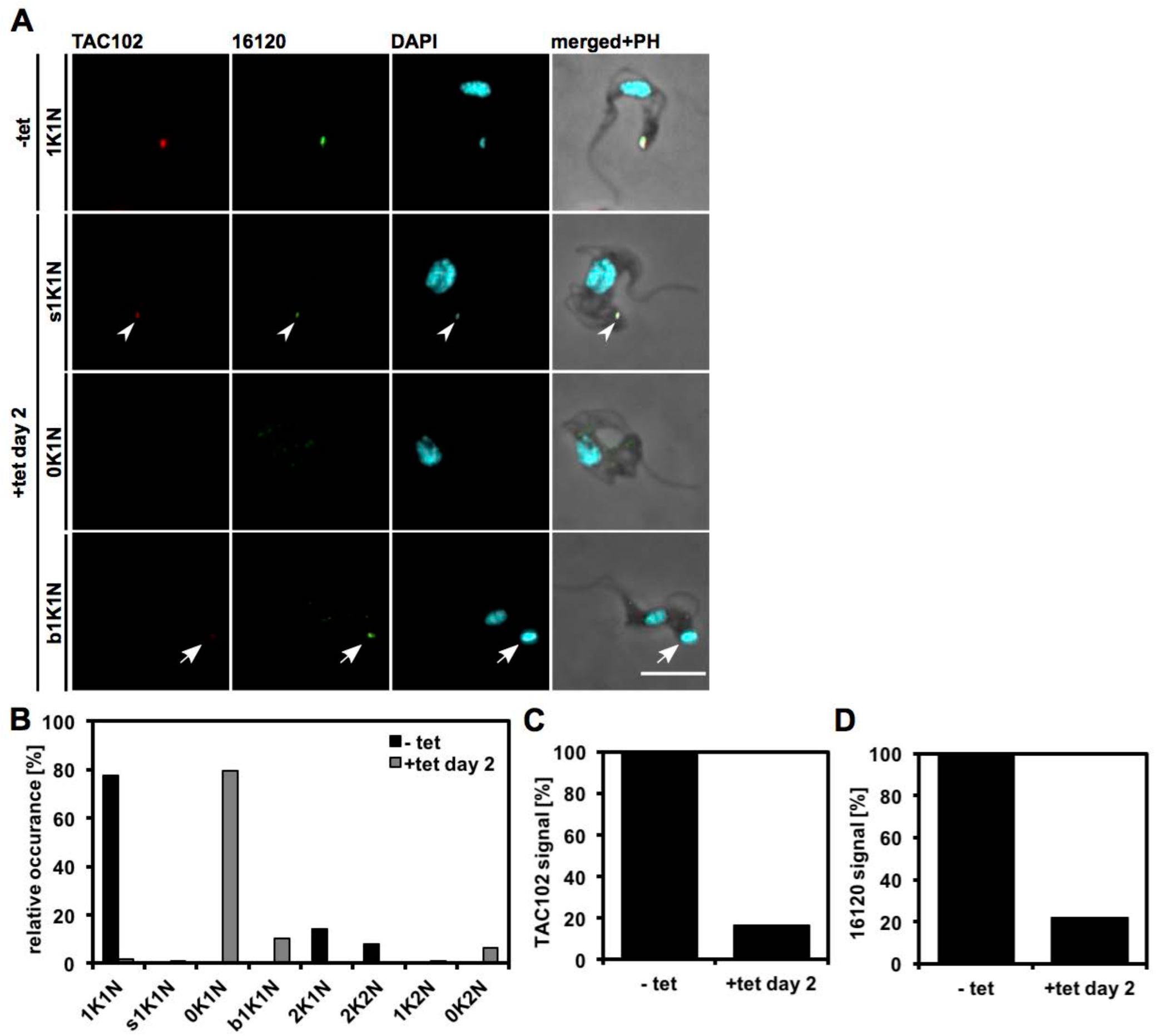
Effect of TAC102 depletion on Tb927.11.16120 in BSF cells. **A)** Immunofluorescence microscopy of TAC102 RNAi Tb927.11.16120-PTP expressing BSF cells. Images were generated as described in the previous figures. **B)** Quantification of the relative occurrence of kDNA discs and nuclei (n≥100 for each condition). **C)** Quantification of TAC102 signals in uninduced cells (-tet) and cells at day three post induction (+tet days 2). **D)** Quantification of Tb927.11.16120-PTP signals in uninduced cells (-tet) and cells at day three post induction (+tet days 2). bK, big kDNA; K, kDNA; N, nucleus; PH, phase contrast; sK; small kDNA. Scale bar: 5 μm.

### Recombinant Tb927.11.16120 binds to DNA without sequence specificity

Due to their vicinity to the kDNA, we wanted to the test whether Tb927.11.16120 and/or TbmtHMG44 bind to DNA. For this, we performed electromobility shift assays (EMSAs) using a 73bp stretch of the conserved minicircle region as a bait and purified recombinant MBP-TbmtHMG44 or Tb927.11.16120-His protein. We detected a shift of the DNA to the top of the gel when 200 ng of Tb927.11.16120-His was added to a reaction containing 20 fmol of DNA bait (Figure 10A). By adding 200-fold excess of the same non-biotinylated DNA the shift could be reversed. To determine the minimal amount of Tb927.11.16120-His needed to obtain a shift of the DNA, we performed a titration with 200 ng, 100 ng, 50 ng, 25 ng and 6.25 ng. With 200 ng and 100 ng of Tb927.11.16120-His almost all DNA was shifted, with 50 ng of Tb927.11.16120-His still more than half of the DNA was shifted, whereas with 25ng less than 50% of the DNA was shifted (Figure 10A). To test whether Tb927.11.16120-His binds to DNA in a sequence unspecific manner, we also performed an EMSA with Tb927.11.16120-His using the Epstein-Barr Nuclear Antigen (EBNA) control DNA (provided with the Thermo Scientific EMSA kit) as a bait. The control reaction with EBNA extract showed that the EBNA DNA could also be shifted with the conditions applied in our experiments (Figure 10B, left side). When using Tb927.11.16120-His we also detected a shift of the EBNA DNA (Figure 10B, right side). From this we can conclude that Tb927.11.16120-His binds to double stranded DNA in a sequence unspecific manner. Furthermore the protein was also able to bind to an RNA oligomer and single stranded DNA, although the shift was weaker and the shifted complex migrated faster. (Figure 10C). We did not observe any DNA binding activity for TbmtHMG44 under the conditions tested (Figure S12). In summary Tb927.11.16120-His seems to bind to DNA and RNA oligomers, while the HMG box containing protein TbmtHMG44 does not.

**Figure 10.**
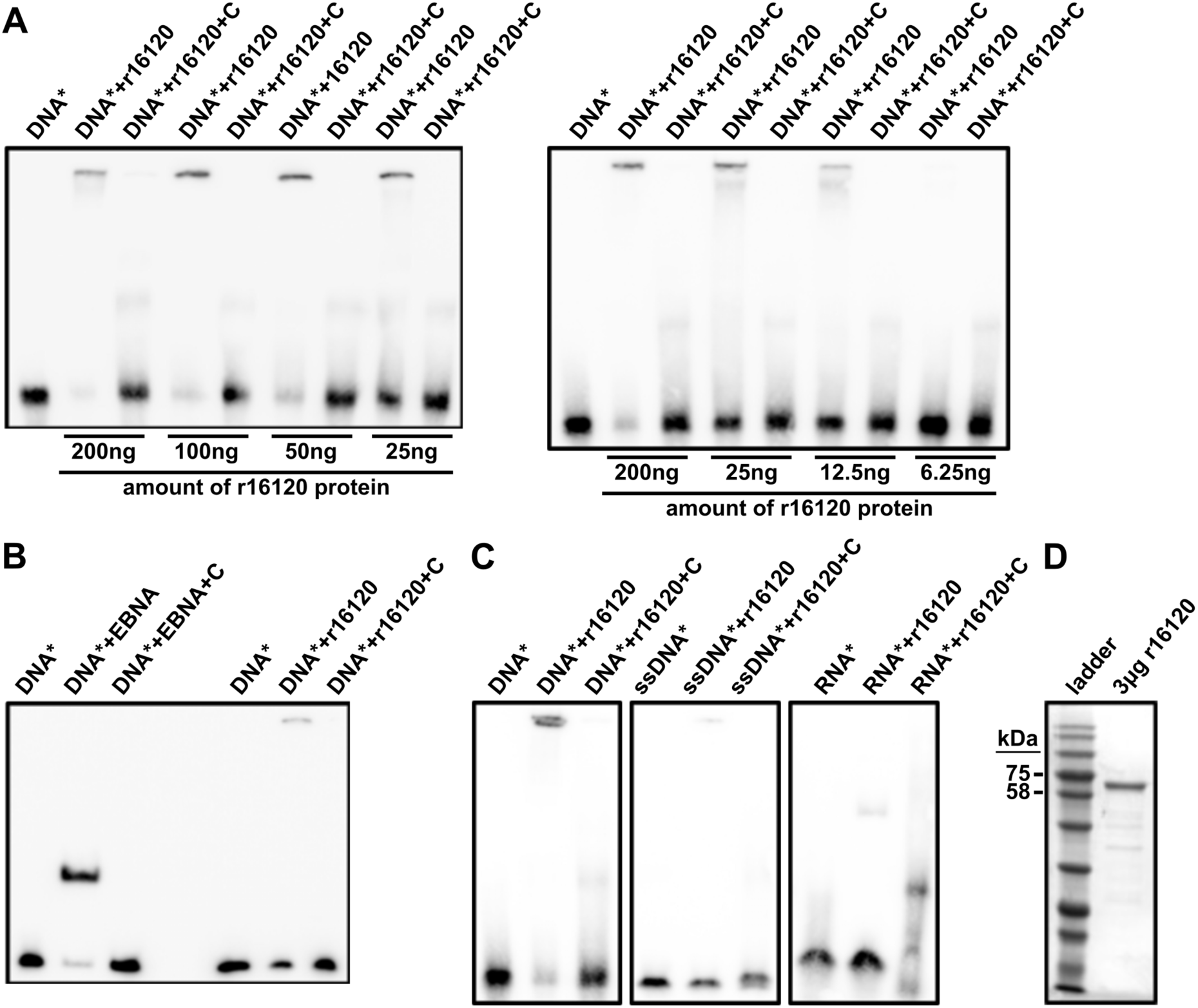
Analysis of the Tb927.11.16120 nucleic acid binding activity using electro mobility shift assays (EMSA). **A)** Titration of purified recombinant Tb927.11.16120-His protein (r16120) to determine minimum amount of protein needed to shift DNA. We used a biotinylated conserved minicircle DNA sequence as a bait. 200-fold excess of the same but non-biotinylated DNA were used as competitor. **B)** Left side: EMSA using biotinylated EBNA DNA and an EBNA nuclear protein extract. 200-fold excess of the same DNA, but non-biotinylated, was used as competitor. Right side: EMSA using the same EBNA DNA but 200ng purified r16120 protein. **C)** EMSAs using either single-stranded DNA or RNA as a bait. 200ng of r16120 were used. **D)** Coomassie stained gel showing purified recombinant r16120. c, 200-fold excess of non-biotinylated competitor DNA; DNA*, biotinylated DNA; r16120, purified recombinant Tb927.11.16120-His.

## Discussion

The depletomics approach confirmed our current model of the hierarchical organization of the TAC since none of the known TAC components except of p166 have been depleted upon TAC102 RNAi. The decrease in abundance of p166 during TAC102 RNAi that has been described before (Hoffmann et al., 2018) and is likely due to the direct interaction of the two proteins which has recently been discovered (Baudouin et al., 2019). The depletomics further showed that a significant number of kDNA replication factors co-fractionated with the isolated, DNase treated flagella, supporting the idea that the mitochondrial genome segregation and replication machinery are linked (Amodeo et al., 2018). The replication proteins identified include the essential replicative mini- and maxicircle polymerases POLIB and POLIC that localize to the KFZ and PolID that displays a dynamic localization between the KFZ and the antipodal sites (see Table S2). The repair polymerase PolIA, which is mostly localized in the mitochondrial matrix was not detected. Surprisingly we also found several proteins that have been described to be localized inside the kDNA disc including the gap repair polymerase POL beta PAK and the HMG-box containing proteins KAP3, KAP4 and KAP6 that are thought to fulfill the role of histone proteins in the kDNA (Wang et al., 2014; Xu et al., 1996). Also the third compartment, the antipodal sites, seem to be linked to the TAC even without the kDNA being present. As representative proteins of antipodal sites we detected the two topoisomerases (TOPIa and TOP2), p93, one helicases (Pif5) as well as the gap filling polymerase POL beta and the replication factor MiRF172. At this point we cannot exclude that some kDNA, which was protected from nucleolytic cleavage remained in the depletomics samples and retained proteins attached to it. However, we consider it unlikely since in that case we should have seen significant differences between the wild type and RNAi induced samples from all three experiments; after all, three days post induction of RNAi targeting any of the three proteins, the vast majority of the cells had strongly reduced amounts of kDNA or lost it all together (see Figure 5, S8 and (Trikin et al., 2016)). Overall, the depletomics data suggests that the kDNA replication machinery is assembled in several different compartments around the kDNA. Some but not all components of the replication machinery depend on the presence of the TAC and many of them seem not to depend on the presence of the kDNA.

The localization of TbmtHMG44 and Tb927.11.16120 between TAC102 and the kDNA and their distribution pattern during kDNA replication is consistent with that of a TAC component (Figure 2, 3, S6), (Hoffmann et al., 2018)). However, although the functional studies by RNAi show that both proteins are required for kDNA maintenance, their depletion does not lead to the characteristic missegregation phenotype seen for other TAC components (Schneider and Ochsenreiter, 2018). Furthermore, different from a typical TAC component the localization of TbmtHMG44 and Tb927.11.16120 depends on the presence of kDNA prior to assembly of the proteins at the TAC kDNA interface (Figure 4, S7) suggesting a more direct interaction with the mitochondrial genome. Interestingly, once TbmtHMG44 and Tb927.11.16120 are assembled at the TAC kDNA interface, the kDNA is no longer required and can be removed by nuclease treatment from isolated flagella (Figure 3, S6).

We analyzed the function of both proteins *in vivo* and *in vitro* and could show that they are required for maintenance of the kDNA and that Tb927.11.16120 binds to DNA. Depletion of Tb927.11.16120 leads to a rapid loss of mini- and maxicircles (Figure 6). Also, the replication intermediates (covalently closed minicircles and the nicked/gapped minicircles) continue to decline at a similar rate (Figure 6), which is different from replication factors like for example PolIB, PolIC, p38, Pri2 or MiRF172, where certain replication intermediates accumulate and eventually lead to the decrease in kDNA (Amodeo et al., 2018; Bruhn et al., 2010; Hines and Ray, 2011; Klingbeil et al., 2002; Liu et al., 2006). Thus both proteins are very likely not directly involved in the replication of DNA. Interestingly, similar to the Tb927.11.16120 RNAi phenotype, TbmtHMG44 RNAi leads to a loss of minicircles including the replication intermediates (Figure S10), however at the same time the maxicircles increase in abundance (Figure S10), a phenomenon that has also been observed during depletion of TopoIA an enzyme responsible for processing minicircle replication intermediates (Scocca and Shapiro, 2007). One could speculate that the loss of the minicircle specific primase Pri2, together with some of the other factors depleted during TbmtHMG44 RNAi allow parts of the replication machinery to be redirected to the maxicircles, thus increasing the replication of this DNA species. Interestingly, aside from the minicircle specific primase Pri2 none of the replication factors that co-fractionated with isolated flagella are decreased in abundance upon Tb927.11.16120 or TbmtHMG44 RNAi (Table S2).

Tb927.11.16120 but not TbmtHMG44 seems to bind to DNA as we demonstrated *in vitro* by EMSA, however we have been unable to show any sequence specificity. The lack of DNA binding specificity might be due to the fact that posttranslational modifications or other interacting proteins are missing. Alternatively, non-specific binding to the minicircles or maxicircles might be the mode of interaction.

Using electron microscopy in combination with differential staining techniques the ULF region of the TAC was previously divided into the inner and outer ULF. The outer ULF consist of acidic proteins, while the inner region contains DNA and mostly basic proteins (Gluenz et al., 2007). How does this fit the data presented here? We now know that the acidic TAC/ULF protein p166 is associated with the inner mitochondrial membrane and directly interacts with the basic more kDNA proximal ULF protein TAC102 (Baudouin et al., 2019; Trikin et al., 2016; Zhao et al., 2008). Towards the kDNA TAC102 is connected to several proteins including the recently characterized basic protein TAP110 (pI 8, (Amodeo et al., 2020)) which is closer to the kDNA than TAC102. We have no evidence that TAC102 binds directly to DNA and based on recent U-ExM data there is clearly a gap between TAC102 and the DNA supporting the model that there are additional proteins between TAC102 and the kDNA (Amodeo et al., 2020). Two of these proteins are likely Tb927.11.16120 and TbmtHMG44, which are both very basic proteins and localize potentially closer to the kDNA than TAC102. For Tb927.11.16120 we could clearly demonstrate DNA binding in vitro making it one of the best candidates to physically connect the kDNA and the TAC.

## Materials and Methods for the analysis of TbmtHMG44

### *T. brucei* cell culture conditions

Monomorphic New York single marker (NYsm) bloodstream form (BSF) *T. brucei* cells (Wirtz et al., 1999) and single marker γL262P T. brucei cells (Dean et al., 2013) were cultured in Hirumi-modified Iscove’s medium 9 (HMI-supplemented with 10% fetal calf serum (FCS) (Hirumi and Hirumi, 1989) and 2.5 μg/ml, geneticin at 37°C and 5% CO_2_. 29-13 double marker procyclic form *T. brucei* cells (Wirtz et al., 1999) were cultured in semi-defined medium-79 (SDM-79) supplemented with 10% FCS, 15 μg/ml geneticin and 25 μg/ml hygromycin at 27°C. For the analysis of the RNAi phenotype, cells containing RNAi constructs, were grown with 1 μg/ml tetracycline or without tetracycline and kept in the exponential phase. For the growth curves we measured the cell density by using a Neubauer chamber to count cell density every 24h. After every counting, the cells were diluted to 10^5^ cells/ml. The BSF single marker and PCF double marker cell lines were obtained from the established collection of the Institute of Cell Biology, University of Bern, Bern, Switzerland. The single marker γL262P cell line is a kind gift of Achim Schnaufer. The NYsm TAC102 RNAi and the single marker γL262P p197 RNAi cell lines were obtained from earlier studies (Hoffmann et al., 2018; Trikin et al., 2016). Depending on the cell line 2.5 µg/ml geneticin, 0.5 µg/ml puromycin, 2.5 µg/ml phleomycin, 5 µg/ml blasticidin or 2.5 µg/ml hygromycin was added to the media. For inducing RNAi 1 μg/ml tetracycline was used.

### YFP-TAC102 immunoprecipitation (IP) and mass spectrometry analysis of the IP

TAC102 was N-terminally YFP-tagged in SmOxP927 cells by using the pPOTv4 vector as previously described (Dean et al., 2015). We used the following primers for the tagging: FWD 5’-AAAGAGTGAGTGAGG-TGAGAGCGAAGAATTGCGGACAGCGCACTTCATACTCTGATCTTTCCCTTTACCCTAGCGACAAAGTATAATGCAGACCTGCTGC-3’, REV 5’-TGAGAGCCAGAGTGGTCAGCCTTCCTTGAAGCAGCGGATTCCTTCCGATCCTGCTTAGCGCCGCACGAGGCCGATACATACTACCCG-ATCCTGATCC-3’. The transfection was achieved by electroporation (1.3kV for 100 μs). The cell preparation and mass spectrometry were performed as previously described (Varga et al., 2017). In brief: 10^10^ cells were cytoskeleton extracted and sonicated (3×10 sec at 10 microns amplitude) to break cellular microtubules, followed by a 30 min incubation with PEME, 0.2 M NaCl and Protease Inhibitors at 4°C. After centrifugation the pellet was resuspended in 0.05% Nonidet P-40 in PBS, and was further fragmented by sonication (6×20 sec at 10 microns amplitude) and added to dynabeads (Thermo Fisher Scientific) crosslinked with the α-GFP antibody (Roche). Beads were incubated with 50 mM Tris pH 7.5, 0.3% SDS and 1mM EDTA to elute the bound material. The eluted sample was then fractionated by SDS-polyacrylamide gel electrophoresis and visualized by Sypro Ruby. Each lane was cut into small pieces and send to mass spectrometry, omitting the tubulin band. The rest was analyzed with the Q Exactive mass spectrometer. MS/MS spectra were searched against a database based on *T. brucei* genome version 9.0 using the Central Proteomics Facility Pipeline version 2.1 of Sir William Dunn School of Pathology, University of Oxford. The enrichment of the proteins was calculated by the spectral index ratio of the eluate to the flow-through. The abundance represents the spectral index ratio of the eluate to the median of the eluate.

### Flagellar extraction for fluorescence microscopy and mass spectrometry analysis

We used five million cells for flagellar extraction for fluorescence microscopy and 20 million cells for flagellar extraction for mass spectrometry analysis. The cell culture was supplemented with EDTA (pH 8.0) to a final concentration of 5 mM prior to centrifugation at 2’500 rcf for 8 min. The pellet of cells was then washed with 1ml of extraction buffer basic (10 mM NaH_2_PO_4_, 150 mM NaCl, 1 mM MgCl_2_ at pH 7.2). Then cells were resuspended in extraction buffer I (extraction buffer basic containing 0.5% Triton X-100; resuspension ratio: 5million/20μl) and extracted for 10 min on ice. The extracted cells were then collected by centrifugation at 3’000 rcf for 3 min at 4°C and washed with extraction buffer basic. For depolymerization of subpellicular microtubules the extracted cells were resuspended in extraction buffer II (extraction buffer basic containing 1 mM CaCl_2_; resuspension ratio: 5million/30μl) and incubated for 45 min on ice. For DNaseI treatment the extraction buffer II was supplemented with DNaseI (Roche) to a final concentration of 100 μg/ml prior to resuspension. The flagella were then collected by centrifugation at 3’000 rcf for 3 min at 4°C and washed twice with PBS. All extraction buffers used for the isolation of flagella for mass spectrometry were supplemented with 2x concentrated cOmplete protease inhibitor cocktail (Roche). Tb927.11.16120 flagellar extraction for immunofluorescence microscopy was performed as described above. For the TbmtHMG44 flagellar extraction, cells were cytoskeleton extracted with 1% Nonidet P-40 in 100 mM PIPES pH 6.9, 1 mM MgSO_4,_ 100 mM EDTA and 2 mM EGTA for 5 min at room temperature. The cytoskeletons were separated from soluble material by centrifugation and then the cytoskeleton pellet was resuspended in 20 mM PIPES pH 6.9 containing 65 mM CaCl_2_ and incubated for 25 min on ice. Following, the flagella were treated either with or without DNase I in DNase buffer (NEB) for 10 min at room temperature. After an additional centrifugation step the pellet was distributed on microscopic slides and the immunofluorescence staining was performed as described further down in this chapter.

### Mass spectrometry and data analysis of flagella

Flagella were extracted as described above from TAC102 RNAi, TbmtHMG44 RNAi and Tb927.11.16120 RNAi cell lines, either non-induced or induced for three days. The isolated flagella were then resuspended in LDS sample buffer (Invitrogen, NU PAGE) and proteins were denatured at 70°C for 10 minutes. The sample preparation and mass spectrometry were performed as previously described (Amodeo et al., 2020). In brief: The protein lysates were each separated on 10% gradient SDS gels (Thermo Scientific) for 8 min at 180 V. Then the proteins were fixed and stained with a Coomassie solution, and the gel lane was cut into slices, minced, and destained. Proteins were reduced in 10mM DTT for 1h at 56°C and then alkylated with 50mM iodoacetamide for 45 min at room temperature in the dark. To obtain peptides, the proteins were digested with trypsin overnight at 37°C and the peptides were extracted from the gel using twice using acetonitrile and ammoniumbicarbonate (Rappsilber et al., 2007). For mass spectrometric analysis, peptides were separated on a 50 cm self-packed column (New Objective) with 75 µm inner diameter filled with ReproSil-Pur 120 C18-AQ (Dr. Maisch GmbH) mounted to an Easy-nLC 1200 (Thermo Fisher) and sprayed online into an Orbitrap Exploris 480 mass spectrometer (Thermo Fisher). We used a 103-min gradient from 3% to 40% acetonitrile with 0.1% formic acid at a flow of 250 nL/min. The mass spectrometer was operated with a top 20 MS/MS data-dependent acquisition scheme per MS full scan. Mass spectrometry raw data were searched using the Andromeda search engine (Cox et al., 2011) integrated into MaxQuant software suite 1.6.5.0 (Cox and Mann, 2008) using the TriTrypDB-46_TbruceiTREU927_AnnotatedProteins protein database (11,203 entries). For the analysis, carbamidomethylation at cysteine was set as fixed modification while methionine oxidation and protein N-acetylation were considered as variable modifications. Match between run option was activated.

### Bioinformatics analysis

Contaminants, reverse database hits, protein groups only identified by site, and protein groups with less than 2 peptides (at least one of them classified as unique) were removed by filtering from the MaxQuant proteinGroups file. Missing values were imputed by shifting a beta distribution obtained from the LFQ intensity values to the limit of quantitation. Further analysis and graphical representation was done in the R framework incorporating ggplot2 package in-house R scripts (R Development Core Team, 2014; Wickham, 2016).

### Extraction of DNA from NYsm and γL262P cells for PCR and Southern blot analysis

Total DNA was isolated from mid-log phase cells. For this, the 50 million cells were centrifuged for 8 min at 2’500 rcf, washed with NTE buffer and then resuspended in 0.5ml NTE buffer. Cells were lysed by addition of 25 μl of 10% SDS. RNA was degraded by addition of 5μl RNase A (20 mg/ml, Sigma) and incubation for 1 h at 37°C. The proteins we degraded by addition of 25 μl proteinase K (10 mg/ml) and further incubation for 2 h at 37°C. For the isolation of DNA 0.5 ml phenol was added to the reaction and mixture was well mixed. Separation of the aqueous and organic phase was performed by centrifugation for 8 min at 16’000 rcf. The aqueous phase was transferred to a new tube and equal amount of chloroform was added mixture was well mixed. The centrifugation was repeated, and the upper phase was transferred to a new tube. For precipitation of the DNA we added 1/10 volume of 3 M Na-acetate (pH 5.2) and 2x the volume absolute ethanol. Then we incubated for 1 h at −80° C or overnight at −20°C. The precipitated DNA was collected by centrifugation at 16’000 rcf for 30 min at 4°C. The pellet was then washed with 1 ml of 70% ethanol and the centrifugation was repeated. After the wash we dried the DNA at 55°C and then resuspended in 50 to 100 μl of sterile MilliQ water.

### Cloning of the tagging and RNAi constructs

For the in situ tagging of Tb927.11.16120 at the C-terminus we used a PTP tag that was integrated into a pLEW100 based plasmid (Schimanski et al., 2005). We obtained the tagging construct by amplification of the Tb927.11.16120 gene ORF positions 1330 to 1806 from genomic NYsm DNA by PCR. For this, we used primers that contained adaptor sequences with the respective restriction sites (FWD 5’-gtacGGGCCCttgtctagtcccatttgggtgactcc-3’, REV 5’-gtacCGGCCGgagtgtggtgccctggggtcttgtg-3’). The amplified ORF region was then cloned into the plasmid by using the ApaI and EagI sites of the plasmid. The resulting tagging plasmid we then linearized with AatII prior to transfection. For the C-terminal HA-tagging of TbmtHMG44 we used the pMOTag4H vector (Oberholzer et al., 2006). We utilized following primers: FWD 5’-TGATGTTCTGGAGCGGACGGGCTGTTTCCGCAGCAAAGAAGCTAACCAGCTGCTT-AGGGAGACGTACATAAACCCTACAAGCAAGAAAAAAGGGAAAGAAGGTaccGGGcccCCCctcGAG-3’ and RV 5’-AAGTAACATAATGCAAC-AAGAAAAGGAGGAAAACATAAAAAGTAATCATGAGAGAGGGAAAAAATGAGAGGAAATGGTTTATGTATCTATAATCGTTACTTGGCGGCCGCTCTAGAACTAGTGGAT-3’. For the cloning of the Tb927.11.16120 RNAi construct we used the ORF position 467 to 966 of the Tb927.11.16120 gene as target. We amplified this ORF region as described above using the following primers: FWD 5’-gtaaGGATCCAAGCTTaggtaaaccggaaggacgtt-3’, REV 5’-cttaCTCGAGTCTAGAatccctgacgttgacgaagt-3’. The TbmtHMG44 RNAi was targeted against the ORF (285 bp – 797 bp; FWD 5’-CTTAAAGCTTGGATCCAGTAACGATTATA-GATACATAAACC-3’ and RV 5’-CTTATCTAGACTCGAG CCGTCACAATCTGCTTCTAC-3’) or against the 3’UTR (1187 bp – 1524bp; FWD 5’-AGTCGAAGCTTGGATCCACACCGAAAAGGCATTCAAC-3’ and RV 5’-TCCGATCTAGACTCGAGACTGGGCAAATAG- CCGTATG-3’). The amplified construct was then inserted twice into a tetracycline (tet) inducible RNAi vector containing a stuffer sequence (modified pLew100-expression vector) (Bochud-Allemann and Schneider, 2002) to generate the hairpin-loop containing dsRNA coding sequence. The restriction sites BamHI, HindIII, XbaI and XhoI were used to do so. The final plasmid we then linearized with NotI prior to transfection. All restriction enzymes we used, were bought from New England Biolabs. The cloning of the TbmtHMG44 (FWD 5’-gatcAAGCTTatgaggcggtgctgttgtg-ccaaaagc-3’, REV 5’-gttaCTCGAGttctttcccttttttcttgcttgtagggtttatgtac-3’) and deltaHMG (FWD 5’-gatcAAGCTTatgaggcggtgctgttgtgccaa-aagcggaaggcctcagtttctcattgacagtccgcacgttggggccatgagagtgccgaac-3’, REV: same as above) overexpression constructs was performed as described previously (Amodeo et al., 2020) using the primers indicated in the brackets.

### Transfections

To obtain transgenic cell lines, we transfected cells with the constructs described above. We integrated the constructs by making use of the homologous recombination mechanism of the cells. The transfection mixtures were prepared as follows: 10 μg of linearized plasmid were mixed with 3x and 1x transfection buffer (90mM Na-phosphate (pH 7.3), 5 mM KCl, 0.15 mM CaCl_2_, 50 mM HEPES (pH 7.3); (Burkard et al., 2007)) to a total volume of 110 μl 1x concentrated transfection buffer. BSF transfection were performed with 40 million cells and PCF transfection was performed with 100 million cells. Cells were collected by a centrifugation of 2’500 rcf for 8 min, mixed carefully with the transfection mixture and transferred to an Amaxa Nucleofector cuvette. The program X-001 or Z-001 of the Amaxa Nucleofector II was used for BSF transfection and the program X-014 for PCF transfections (Schumann Burkard et al., 2011). After transfection the cells were recovered in the respective medium for 20h. After the recovery we then added the respective antibiotics for selection of integration of the transfected constructs. For the selection of the in situ tagging of Tb927.11.16120 we used 5 μg/ml blasticidin for the BSF cells and 10 μg/ml blasticidin for the PCF cells. For selection of the RNAi construct we used 5 μg/ml blasticidin too.

### Quantitative Real-Time PCR

The extraction of total RNA was performed with RiboZol™(Ameresco). 5 µg of RNA was treated with 0.5 µl DNase I (NEB) for 15 min at 37°C. Purification was performed with 200 mM NaOAc pH 4, phenol, chloroform/isoamylalcohol. To synthesize cDNA the Omniscript® reverse transcription kit (Qiagen) was used and performed as described in the manual. For this 1 µg of the template and random hexamers as primers were incubated for 1 hour at 37°C, followed by heat inactivation for 5 min at 93°C. The MESA Green qPCR MasterMix Plus for SYBR assay (Eurogentec) was used to perform the quantitative Real-Time PCR (qPCR) and the cDNA was mixed as described in the manual. The qPCR was performed with the ABI Prism 7000 Sequence Detection System (Applied Biosystems) and the data were analyzed by using the 7000 System SDS software v1.2 (Applied Biosystems). For TbmtHMG44 we used following primer sequences: FWD 5’-ACCAGCTGCTTAGGGAGACG-3’, RV 5’-GAACACCAGCACTCACCCGT-3’. Normalization of the CT-values was performed to the housekeeping gene α-tubulin (FWD 5’-CGCTATTATTAGAACAGTTTCTGTAC-3’, RV 5’-GTTACCAACCTGGCAACCA-3’) and the uninduced value equals one.

### Northern blot analysis for detection of RNA

For the isolation of RNA, we collected cells, 50 millions of each, non-induced NYsm Tb927.11.16120 RNAi cells and cells at day three post induction of the RNAi by centrifugation at 2’500 rcf for 8 min. The pellet was washed with 1 ml PBS and then cells were lysed by resuspension in 1 ml of TRIZOL (Ambion). 0.2 ml chloroform was added to the lysed cells and the mixture was vortexed for approximately 15 s prior to centrifugation at 1’100 rpm for 10 min at 4°C. The aqueous phase was then transferred to a fresh tube and mixed with the equal volume of isopropanol. Again, the mixture was vortexed for around 15 s prior to centrifugation as described above. The precipitated RNA was then washed with 1 ml of 70% ethanol (same centrifugation as described above). The isolated RNA was then dried at 55°C and resuspended in 50 to 100 μl nuclease free water. For the northern blot analysis, 10 μg of RNA/sample were used to perform gel electrophoresis. For this we mixed the RNA with sample preparation buffer (0.1 mg/ml ethidium bromide, 15% formaldehyde, 0.1 M MOPS, 0.3 M Na-acetate, 0.05 M EDTA (pH 8), bromophenol blue) and incubated for 15 min at 65°C. After cooling down the sample to room temperature the samples were load onto a 1.4% agarose gel and RNA was resolved at 100V for approximately 2 h. As a running buffer we used MOPS (pH 7) containing 5.92% formaldehyde. After the electrophoresis the RNA was transferred to a Hybond nylon membrane by capillary transfer with 20x SSC (3 M NaCl, 0.3 M Na-citrate, pH 7) over night. After the transfer the membrane was auto-crosslinked with the Stratagene UV-Stratalinker. For detection of the Tb927.11.16120 mRNA, membranes were first pre-wet in 5xSSC and then blocked in hybridization solution (5x SSC, 1:12.5 100x Denhardt’s (2% BSA, 2% polyvinylpyrrolidone, 2% Ficoll), 50mM NaHPO_4_ (pH 6.8), 1% SDS, 100 μg/ml salmon sperm DNA) at 65°C for one hour. To generate the probe, we used the same PCR product which we also used for cloning of the RNAi construct. In a first step, the DNA was denatured at 95°C for 5 min. Then we followed the manufacturer’s protocol for radioactive labelling through Klenow (Random primed DNA labeling kit, Roche). The reaction was stopped by the addition of 0.2 M EDTA (pH 8.0) and added to the membrane in hybridization solution. The membrane was hybridized overnight at 65°C. The next day membrane was washed twice with 2x SSC, 0.1% SDS and twice with 0.2x SSC, 0.1% SDS at 60°C. Then the membrane was exposed to a storage phosphor screen (Amersham Bioscience) for around 24h and scanned by a Storm PhosphoImager (Amersham Bioscience). For normalization, the membrane was blocked as described above and then probed for the 18S RNA. The 18S rRNA probe was generate as follows: 1.8 μl 18S rRNA (10 μM) was mixed with 12.5 μl water, 2.7 μl gamma-32P-ATP (1 MBq), 2 μl PNK buffer (10x) and 1 μl T4 PNK and incubated for 30 min at 37°C. The reaction was stopped with 5 μl EDTA (0.2 M) and 75 μl TE buffer and incubated 5 min at 98° C. The probe was then quenched for 2 min on ice and 50 μl of the reaction was added to the membrane in hybridization solution. Hybridization and washes were performed as described above. Exposure was approximately 15 min instead of 24 h and the screen was scanned as described above.

### Immunolabeling for microscopy

Approximately one million cells were collected by centrifugation for 3 min at 1’800 rcf. Cells were washed with 1 ml PBS and then resuspended in 20 μl PBS and spread on a glass slide for regular epifluorescence microscopy or on a plasma coated Nr. 1.5 cover glasses (Marienfeld) for 2D STED microscopy. After the cells were settled, we fixed them for 4min with 4% PFA (in PBS). After fixation we permeabilized with 0.2% Triton-X 100 (in PBS, 5 min). For immunolabeling of flagellar extracts, we resuspended five million flagella as described above and fixed them as described above. Cells and flagella were blocked with 4% BSA in PBS for 30 min. The primary and secondary antibodies were incubated for 45 min to 1h and diluted in blocking solution as follows: polyclonal rabbit-anti-Protein A (Sigma) detecting the PTP epitope 1:2’000, rat YL1/2 antibody detecting tyrosinated tubulin as present in the basal body (Kilmartin et al., 1982) 1:100’000, monoclonal mouse TAC102 antibody (Trikin et al., 2016) 1:2’000, rabbit-anti-HA (Sigma) 1:1’000, rabbit-anti-ATOM40 1:10’000, Alexa Fluor® 488 Goat-anti-Rabbit IgG (H+L) (Invitrogen), Alexa Fluor® 594 Goat-anti-Mouse IgG (H+L) (Molecular probes), Alexa Fluor® 488 Goat-anti-Mouse IgG (H+L) (Invitrogen), Alexa Fluor® 647 Goat-anti-Rat IgG (H+L) (Life technologies), Alexa Fluor® 594 Goat-anti-Rat IgG (H+L) (Invitrogen) all 1:700 or 1:1000. For 2D STED microscopy we used antibodies with following dilutions: Polyclonal rabbit-anti-Protein A antibody (Sigma) and monoclonal mouse-anti-TAC102 antibody as described above and the secondaries Alexa Fluor® 594 goat-anti-Rabbit IgG (H+L) (Invitrogen) and the Fluor® Atto647N Goat-anti-Mouse IgG (H+L) (Molecular probes) we used 1:500 in 4% BSA and incubated for one hour on the cover slips. After each antibody incubation cells were washed 3x with 0.1 % Tween-20 (in PBS). A final wash with with PBS was performed prior to mounting. Cells were mounted with ProLong® Gold Antifade Mounting Medium with DAPI (4’,6-diamidine-2-phenylindole) (Invitrogen). The YL1/2 antibody is a kind gift of Keith Gull. The ATOM40 antibody is a kind gift from André Schneider.

For super-resolution microscopy, cells were spread on a glow discharged cover glas (coverslip thickness no. 1.5, plasma coating for 30 s with FEMTO SCIENCE CUTE discharger) instead of a slide.

### Epifluorescence, confocal and 2-dimensional stimulated emission depletion microscopy (2D STED)

For regular epifluorescence microscopy images were acquired with the Leica DM5500 B microscope with a 100x oil immersion phase contrast objective and analyzed using LAS AF software (Leica) and ImageJ. For confocal and 2D STED super-resolution microscopy we used the SP8 STED microscope (Leica, with a 100× oil immersion objective and the LAS X Leica software). Images were acquired as z-stacks with a z-step size of 120 nm and a X-Y resolution of 37.9 nm. We used the 594 nm excitation laser and the 770 nm depletion laser to acquire the TbmtHMG44 and Tb927.11.16120 signal. The TAC102 signal we acquired with the 647 nm excitation laser and the 770 nm depletion laser. The DAPI signal was acquired with confocal settings. To deconvolute the pictures we used the Huygens professional software.

### Cytoskeleton extraction for fluorescence microscopy

For the cytoskeleton extraction the cells were harvested, washed, resuspended and spread on a slide as described above. Then the cells were incubated for 5min with extraction buffer (100mM PIPES (pH 6.8), 1 mM MgCl2) containing 0.05% Nonidet P-40. After the extraction, the cells were washed twice with extraction buffer and the immunolabeling was performed as described above.

### SDS-PAGE and western blot analysis

Cell extracts were mixed with Laemmli buffer to a final concentration of 5 million cell equivalents/15μl. Either one million or five million cell equivalents (depending on the experiment) were load per lane on an SDS-polyacrylamide gel. The gels were resolved by 80-120V and transferred (wet transfer, 100V for 1h) onto a PVDF Immobilon®-FL transfer membranes (0.45 μm, MILLIPORE), followed by blocking (5% milk in PBS + 0.1% Tween-20 (PBST)) for 1 hour at room temperature. Primary antibodies were incubated for either 1 hour at room temperature or overnight at 4°C. Secondary antibodies were applied for 1 hour at room temperature. All used antibodies were diluted in blocking solution. The following antibodies were used: mouse anti-TAC102 (1:1000,(Trikin et al., 2016)), rabbit anti-HA (1:1000, Sigma), rabbit anti-ATOM40 (1:10000,(Pusnik et al., 2011)), rabbit anti-mouse HRP-conjugated (1:10000, Dako) and swine anti-rabbit HRP-conjugate (1:10000, Dako), rabbit peroxidase anti-peroxidase soluble complex (PAP) (1:2000, for detection of the PTP tag). After antibody incubation, the membranes were washed three times with PBST. A final wash with PBS was performed prior to detection of the protein with the SuperSignal West Femto Maximum Sensitivity Substrate (Thermo Scientific). The AmershamTM Imager 600 (GE Healthcare Life Sciences) was used to visualize the protein bands on the blot.

### Southern blot analysis for the detection of mini- and maxicircle DNA

For the mini- and maxicircles 5 µg of total DNA, digested with HindIII and XbaI, were loaded, while for the free minicircles 8-10 µg of total DNA was used. The experiment was performed as described previously (Amodeo et al., 2018). For detection either the DIG high prime DNA labeling and detection kit (ROCHE) or α-^32^P-dCTP and/or α- ^32^P-dATP were used. Minicircle primers: FWD 5’-TATGGGCGTGCAAAAATACA-3’, RV 5’-CGAAGTACCTCGGACCTCAA-3’. Maxicircle primers: FWD 5’-GCCTGCTGGGAATTGTTCTG-3’, RV 5’-GATCCACGTTCAATGCACCG-3’. Tubulin primers: FWD 5’- CTAACATACCCACATAAGACAG-3’, RV 5’-ACACGACTCAATCAAAGCC-3’.

### Electro mobility shift assay (EMSA)

Encoded by the plasmid pET22b+, C-terminally His-tagged Tb927.11.16120 (Tb927.11.16120-His) and N-terminally MBP-tagged TbmtHMG44 (MBP-5020) were expressed in BL21 *E. coli*. Expression from the corresponding T7 promoter was induced with 1mM isopropyl-β-D-thigalactoside (IPTG) at a growth temperature of 28°C for 4 hours. To purify the recombinant proteins, a amylose resin column (NEB) or a Nickel loaded HisTrap HP column (GE Healthcare) were used. MBP-5020 was purifed accoring to the manufacturer’s protocol and eluted with a buffer containing 20 mM Tris-HCl pH 7.4, 0.2 M NaCl, 1 mM EDTA, 10 mM maltose. For the purification of Tb927.11.16120-His, the cells were harvested by centrifugation, and resuspended on ice in 16ml 20mM Tris, 4mM MgCl_2_ (pH 8 at 4°C) per liter of culture volume. The cells were opened using a Microfluidizer at 10’000 PSI. The lysate was then treated with 500 units of Benzoase Nuclease (Sigma-Aldrich) per liter of culture volume, for 20 min on ice. Triton was added to a final concentration of 0.5% v/v, and the lysate was incubated for 10 min on ice. The lysate was then centrifuged at 3000 rcf for 10 min, and the pellet was resuspended in 83 ml/L 50 mM Tris, 100 mM NaCl, 1 mM EDTA, 10 mM DTT (pH 8 at 4°C), and incubated for 10 min on ice. Insoluble substances were then pelleted at 3000 rcf, for 10 min. The pellet was resuspended in 83 ml/L 50mM Tris, 100 mM NaCl, 1 mM EDTA (pH 8 at 4°C), and incubated for another 10 min on ice. The sample was pelleted at 3000 recf for 10 min. The pellet was resuspended in 16 ml/L 50 mM Tris, 100 mM NaCl (pH 8 at 4°C), and incubated for 10 min. Then it was centrifuged for another 10 min at 3000 rcf and the pelleted inclusion bodies were solubilized in 16 ml/L 8 M urea, 50 mM Tris, 400 mM NaCl (pH 8 at 4°C). Insoluble components were removed with centrifugation, and the recombinant protein was purified on a Nickel loaded HisTrap HP column (GE Healthcare) using the ÄKTA prime system. The protein was eluted in a gradient from 50 mM Tris, 500 mM NaCl to a buffer containing the same ingredients plus 500 mM imidazole, and eluted at about 20% of the gradient, with 100 mM Imidazole. The protein was then dialyzed into 50 mM Tris, 100 mM NaCl overnight, and stored at −20°C. Protein concentrations were measured using the NanoDrop 2000 spectrophotometer. For the MBP-cleavage the AcTEV™ Protease (Invitrogen) was added to the protein as suggested from the manual. The LightShift® Chemiluminescent EMSA assays were performed according to the manufacturer’s kit protocol, with small adaptations in the shift assay (Thermo Scientific). We did not use dI○dC and adapted the type and amounts of detergent as described further down in the text. We used 73 bp or the conserved minicircle sequence as DNA bait: FWD 5’-GAAAAAACGGAAAATCTTATGGGCGTGCAAAAATACACATACACAAATCCCGTGCTTATTTTAGGCCATTTTT-3’ and REV 5’-AAAAATGGCCTAAAATAAGCACGGGATTTGTGTATGTGTATTTTTGCACGCCCATAAGATTTTCCGTTTTTTC-3’. The binding reactions were performed with the kits binding buffer and supplemented with 0.1% digitonin, 0.01% BSA, 20 fmol biotinylated DNA, and 4 pmol unlabeled DNA. Variable amounts of recombinant protein were used. After resolving on a 6% acrylamide gel and transfer onto a nylon membrane, the biotinylated DNA was detected with streptavidin-HRP conjugate according to the manufacturers protocol (Thermo Scientific). As a control the Epstein-Barr nuclear antigen protein extract and DNA were used, as provided with the kit.

### Phylogenetic analysis

Phylogenetic relationship was reconstructed using the Phylo.fr package (Dereeper et al., 2008). For this the sequences were aligned using the MUSCLE algorithm with standard settings (Edgar, 2004). The phylogenetic tree reconstruction was calculated by the PhylML 3.0 algorithm using standard settings (Guindon et al., 2010). Tree visualization as well as the comparison was done using the Phylo.io tool (Robinson et al., 2016).

## Acknowledgments

We thank Keith Gull for the YL1/2 antibody. Imaging was supported by the Microscopy Imaging Center of the University of Bern, Switzerland. The research was supported by grants from the Novartis Foundation and the Swiss National Science Foundation (SNF) (160264). The YFP-TAC102 IP was performed in the lab of Keith Gull with the help of Vladimir Varga. We thank Laura Pfeiffer for her excellent technical assistance, and Nina Suter and Dominic Rothen for their help in generation of some cell lines. Many thanks Caroline Dewar for the RNAi vector.

## Supplementary Figures

**Figure S1:**
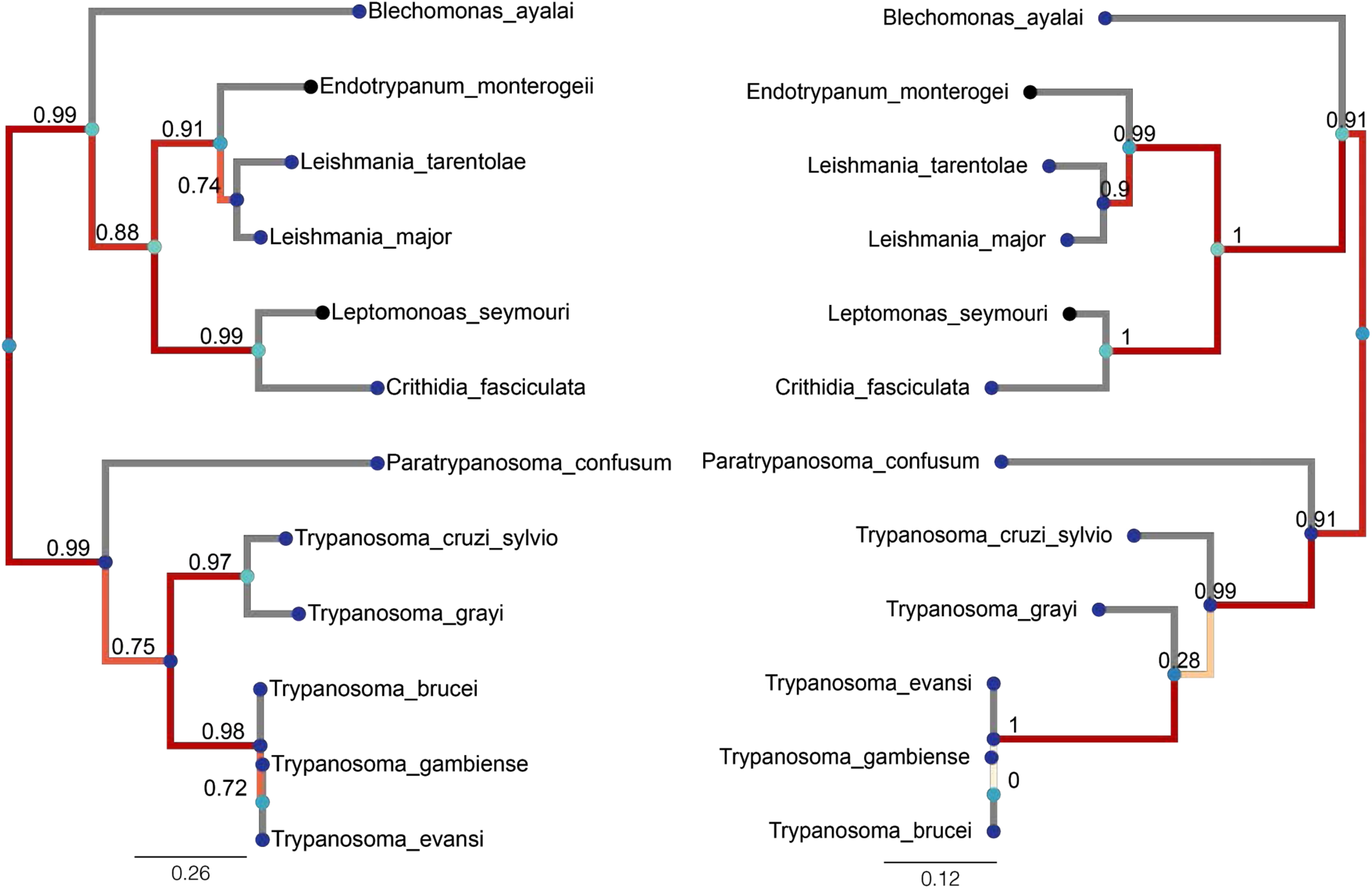
A phylogenetic reconstruction showing the conservation of TbmtHMG44 (left) and Tb927.11.16120 among Kinetoplastids. The scale bar indicates the number of amino acid substitutions.

**Figure S2:**
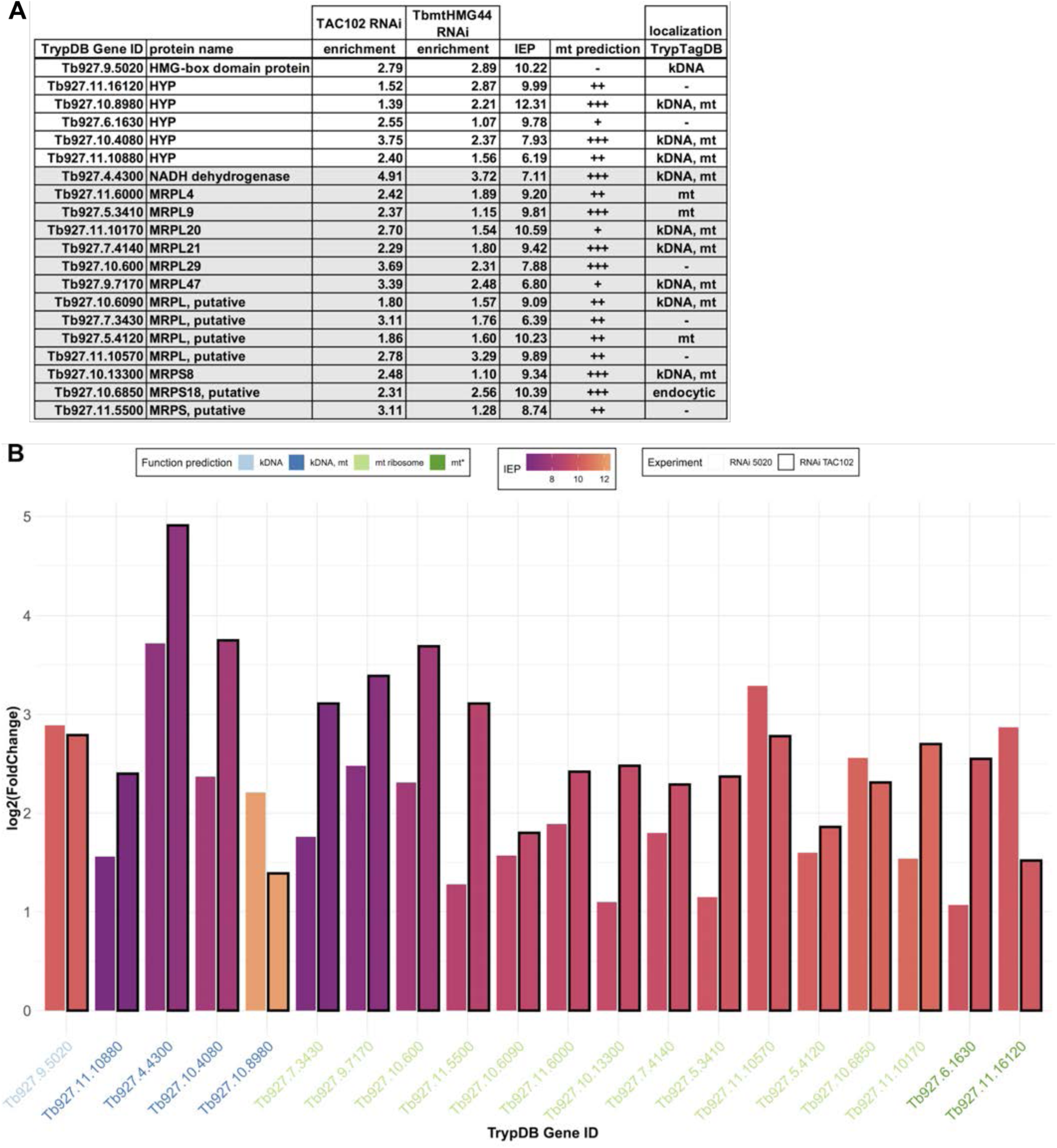
Proteins that significantly changed in expression levels in the TAC102 and TbmtHMG44 RNAi. **A)** Summary of the significantly enriched proteins upon TAC102 and TbmtHMG44 RNAi. Enrichment shows the log_2_ fold-change values values. HYP, hypothetical protein, conserved; IEP, isoelectric point; mt, mitochondrion/mitochondrial; MRPL, mitochondrial ribosomal protein large subunit; MRPS, mitochondrial ribosomal protein small subunit; +, mitochondrial prediction by one of the three data sets/methods: Mitocarta, mitochondrial importome or Mitoprot; ++, mitochondrial prediction by two of the three data sets/methods mentioned before; +++, mitochondrial prediction all three data sets/methods mentioned before. **B)** Visualization of the information above into a barplot. IEP, isoelectric point, mt, mitochondrion; 5020, Tb927.9.5020 (alias TbmtHMG44).

**Figure S3:**
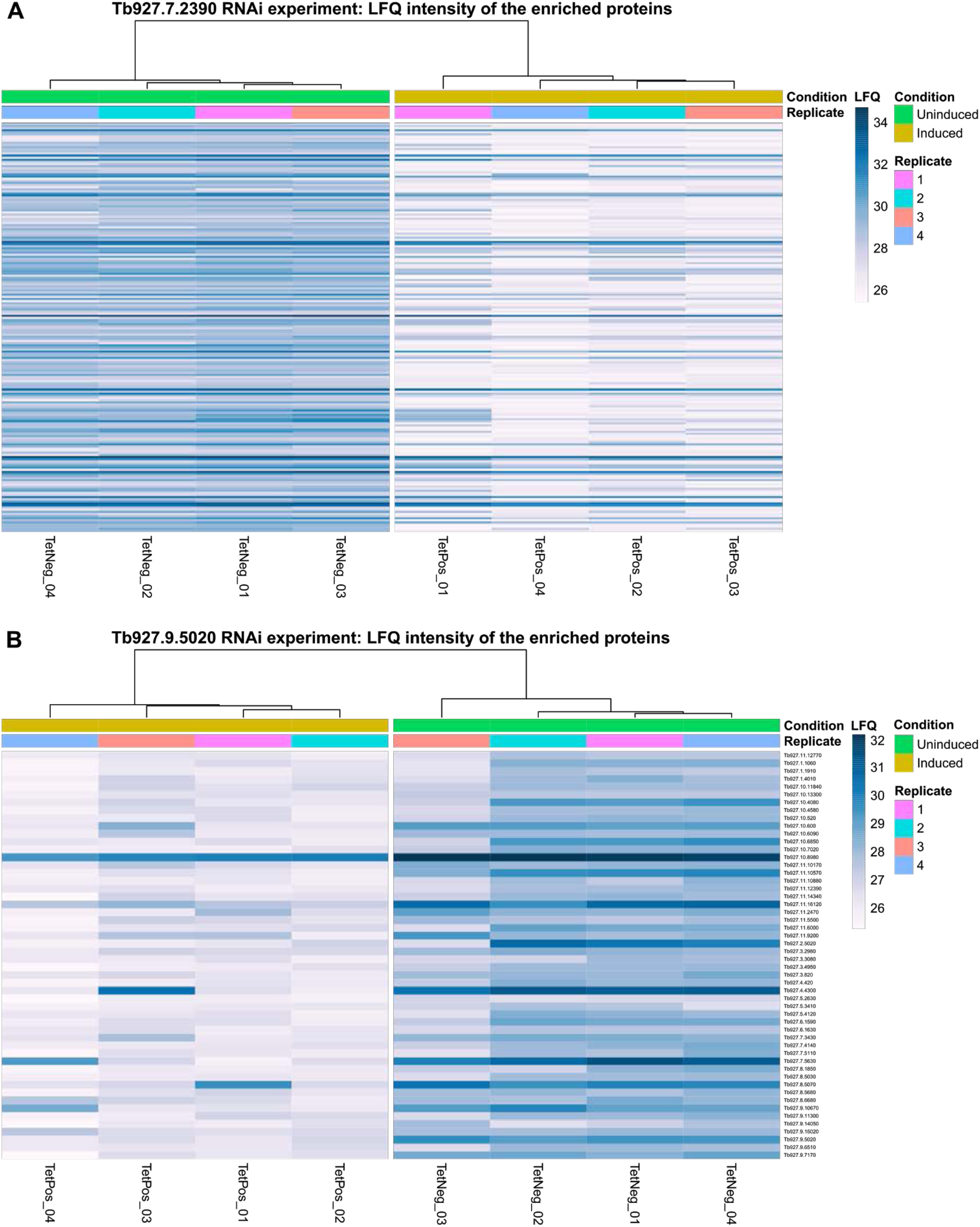
Heat maps of proteins candidates from mass spectrometry analysis of TAC102 and TbmtHMG44 (Tb927.9.5020) RNAi. **A)** Heat map TAC102 RNAi. **B)** Heat map Tb927.9.5020 RNAi.

**Figure S4:**
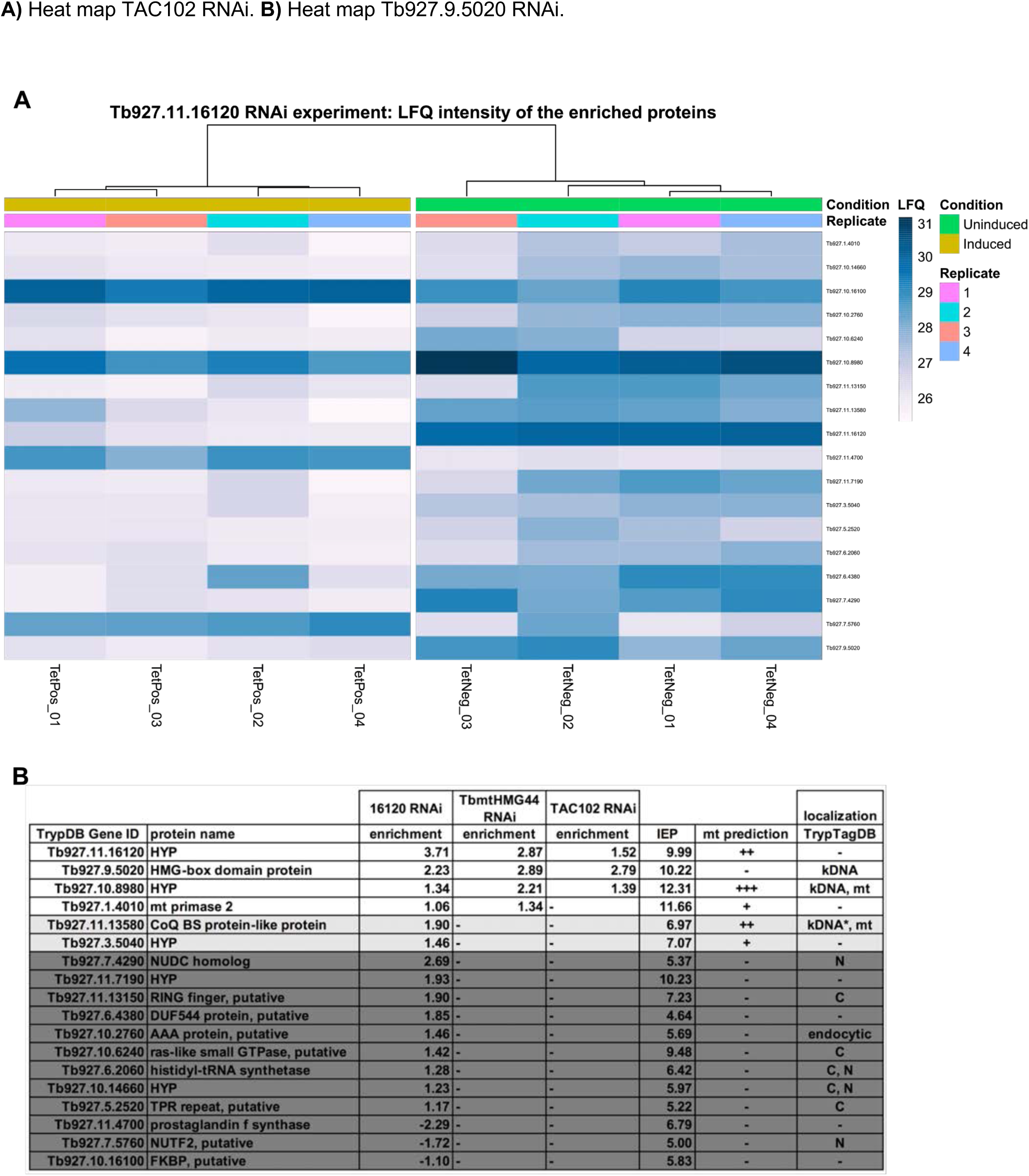
Proteins that depend on the presence of Tb927.11.16120. **A)** Heat map of protein candidates from mass spectrometry analysis of Tb927.11.16120 RNAi. **B)** Summary of the significantly enriched proteins upon Tb927.11.16120 RNAi. Enrichment shows the log_2_ fold-change values. Enrichment shows the log_2_ fold change values. AAA, ATPase family associated; C, cytoplasm; CoQ BS, ubiquinone biosynthesis; DUF, domain of unknown function; FKBP, FK506-binding protein; HYP, hypothetical protein, conserved; IEP, isoelectric point; mt, mitochondrion/mitochondrial; N, nucleus; NUDC, nuclear distribution protein C; NUTF2, nuclear transport factor 2; TPR, tetratricopeptide repeat; +, mitochondrial prediction by one of the three data sets/methods: Mitocarta, mitochondrial importome or Mitoprot; ++, mitochondrial prediction by two of the three data sets/methods mentioned before; +++, mitochondrial prediction all three data sets/methods mentioned before; *, localization cell cycle dependent.

**Figure S5.**
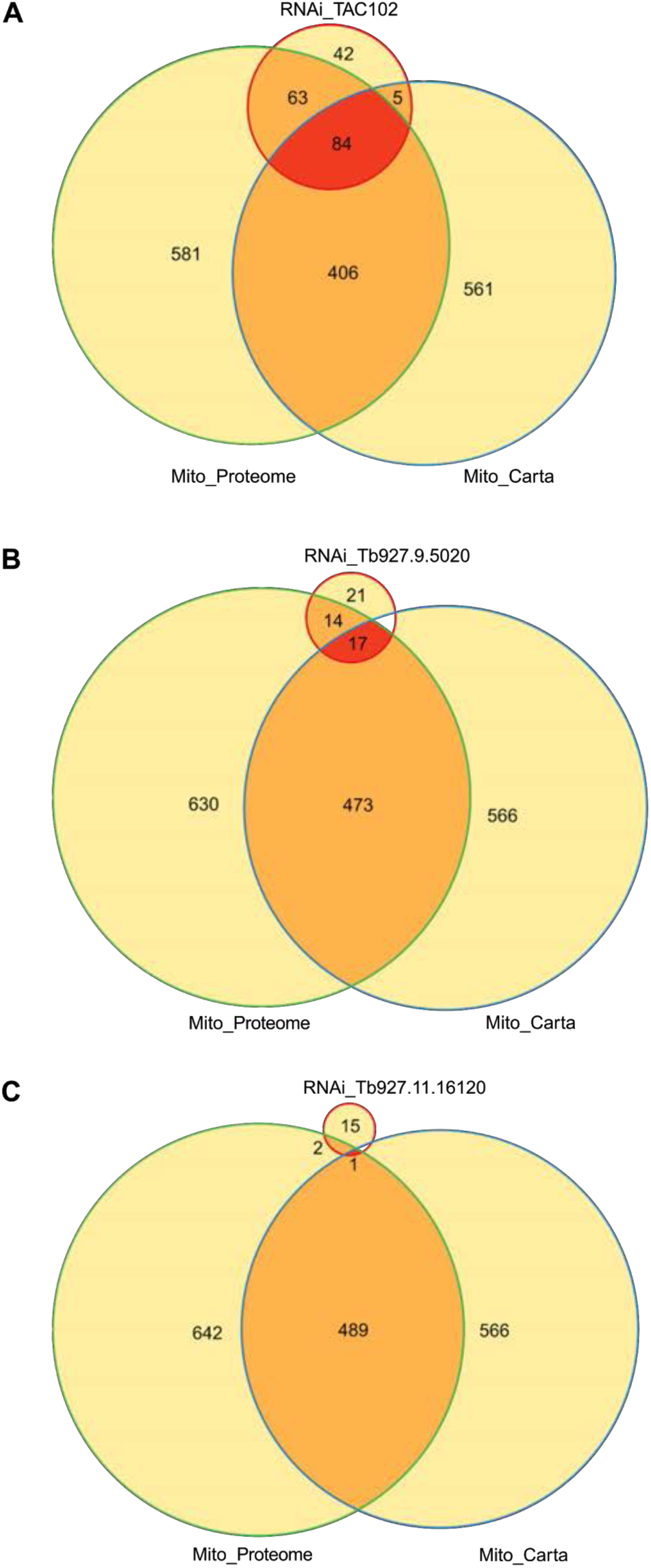
Mitochondrial prediction for the proteins depleted upon TAC102, TbmtHMG44 (Tb927.9.5020) or Tb927.11.16120 RNAi.

**Figure S6.**
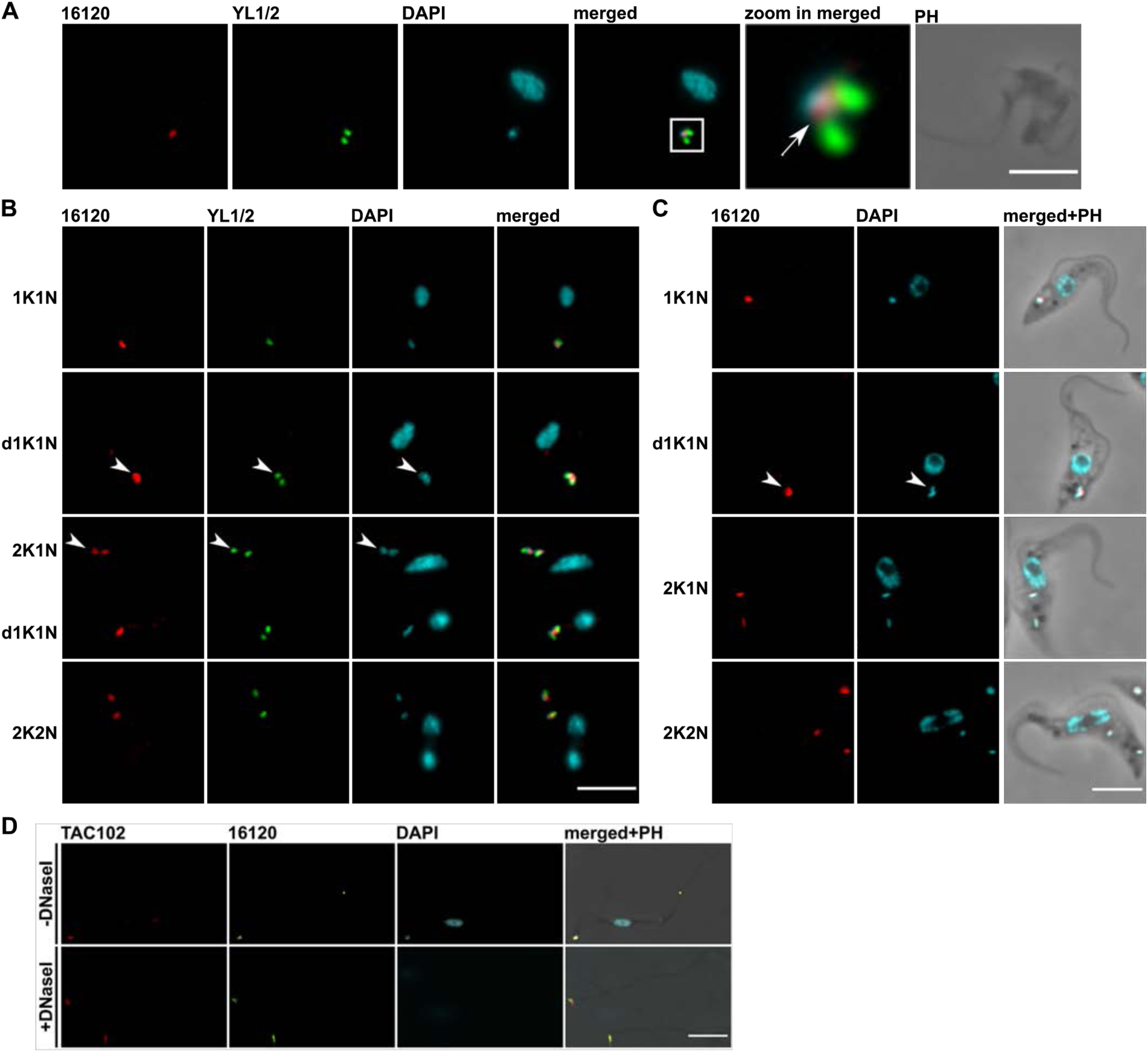
Localization of Tb927.11.16120-PTP in BSF and PCF cells, and isolated flagella. **A)** Representative Immuno-fluorescence microscopy image of a Tb927.11.16120-PTP expressing BSF cell. The signals are represented by maximum intensity projections from image stacks. The mature basal bodies (green) were detected with the YL1/2 monoclonal antibody. Tb927.11.16120-PTP (red) was detected by means of the anti-Protein A antibody. The kDNA and the nucleus were stained with DAPI (cyan). Zoom factor: 5x. The arrow points to the Tb927.11.16120 signal. **B)** The image processing and staining procedure was performed as described in A. Arrowheads point to replicated Tb927.11.16120, basal body and kDNA signals. dK, replicating/duplicated kinetoplast; K, kinetoplast; N, nucleus. Scale bar 5 µm. **C)** The image processing and staining procedure was performed as described in the previous A. dK, replicating/duplicated kinetoplast; K, kinetoplast; N, nucleus; PH, phase contrast. Scale bar, bar 5 µm. **D)** Representative images of isolated flagella either without or with DNaseI treatment during extraction. The image processing and staining procedure were performed as described above. TAC102 (red) was detected with the anti-TAC102 monoclonal antibody. PH, phase contrast. Scale bar, 5 µm.

**Figure S7.**
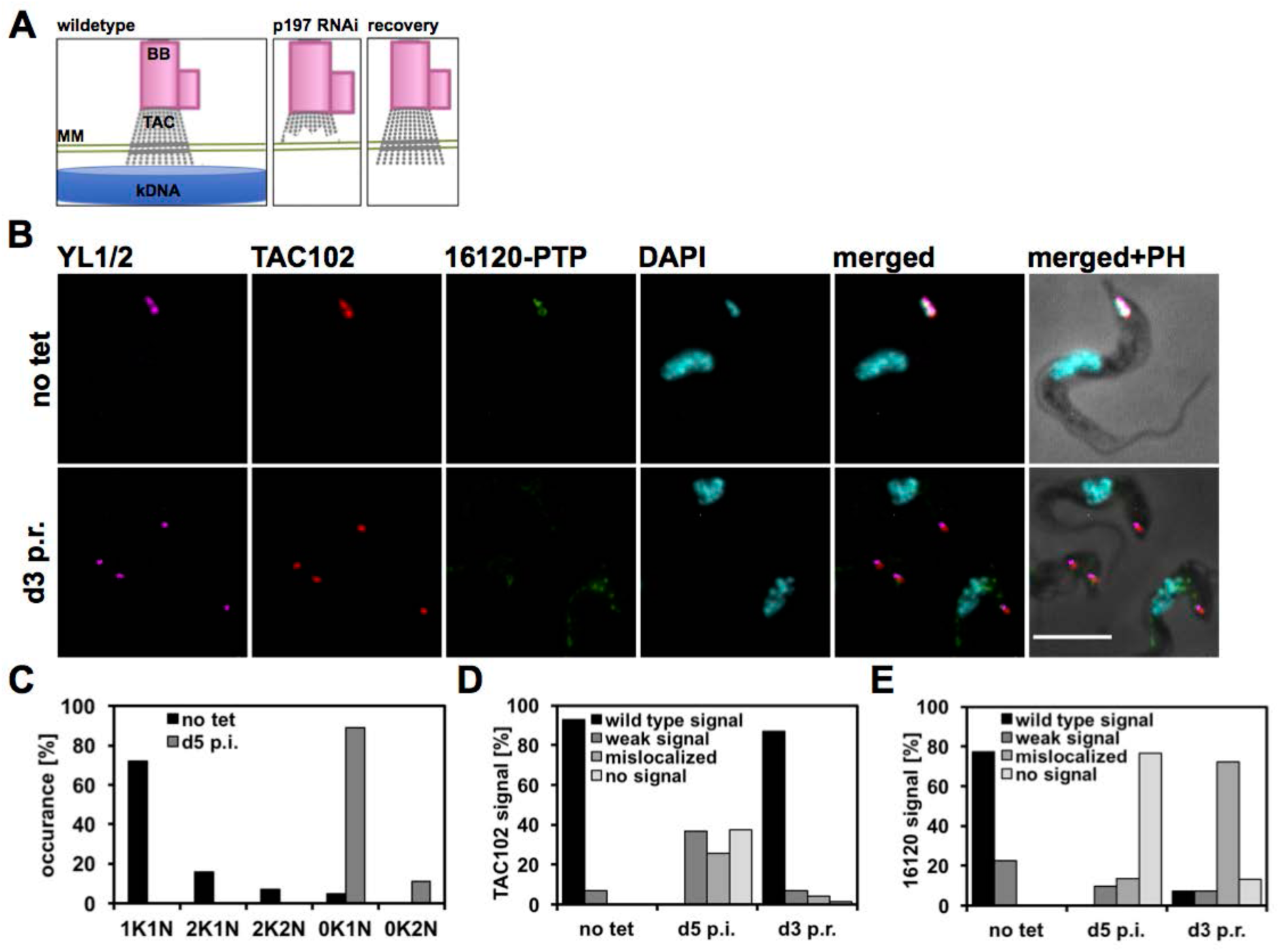
Recovery of Tb927.11.16120 signals in p197 RNAi Tb927.11.16120-PTP expressing γL262P BSF cells. **A)** Mechanism of the p197 RNAi recovery experiment. Non-induced cells contain an intact TAC and kDNA. When p197 RNAi is induced, the TAC gets disrupted. At day five post induction the population is akinetoplastic. When tetracycline is removed, the cells can recover from the RNAi. The TAC gets reassembled without kDNA. **B)** Immunofluorescence microscopy of the cell line described above. Staining procedure, image acquiring and processing was performed as described in previous figures. Non-induced (no tet) and cells at day three post recovery (d3 p.r.) are shown. **C)** Quantification of the relative occurrence of kDNA networks and nuclei (n≥100 for each condition). **D)** Quantification of TAC102 signals (n≥100 for each condition). **E)** Quantification of Tb927.11.16120 signals (n≥150 for each time point). BB, basal body; d5 p.i., induced cells at day five post; K, kDNA; N, nucleus, PH, phase contrast; TAC, tripartite attachment complex. Scale bar: 5 μm.

**Figure S8:**
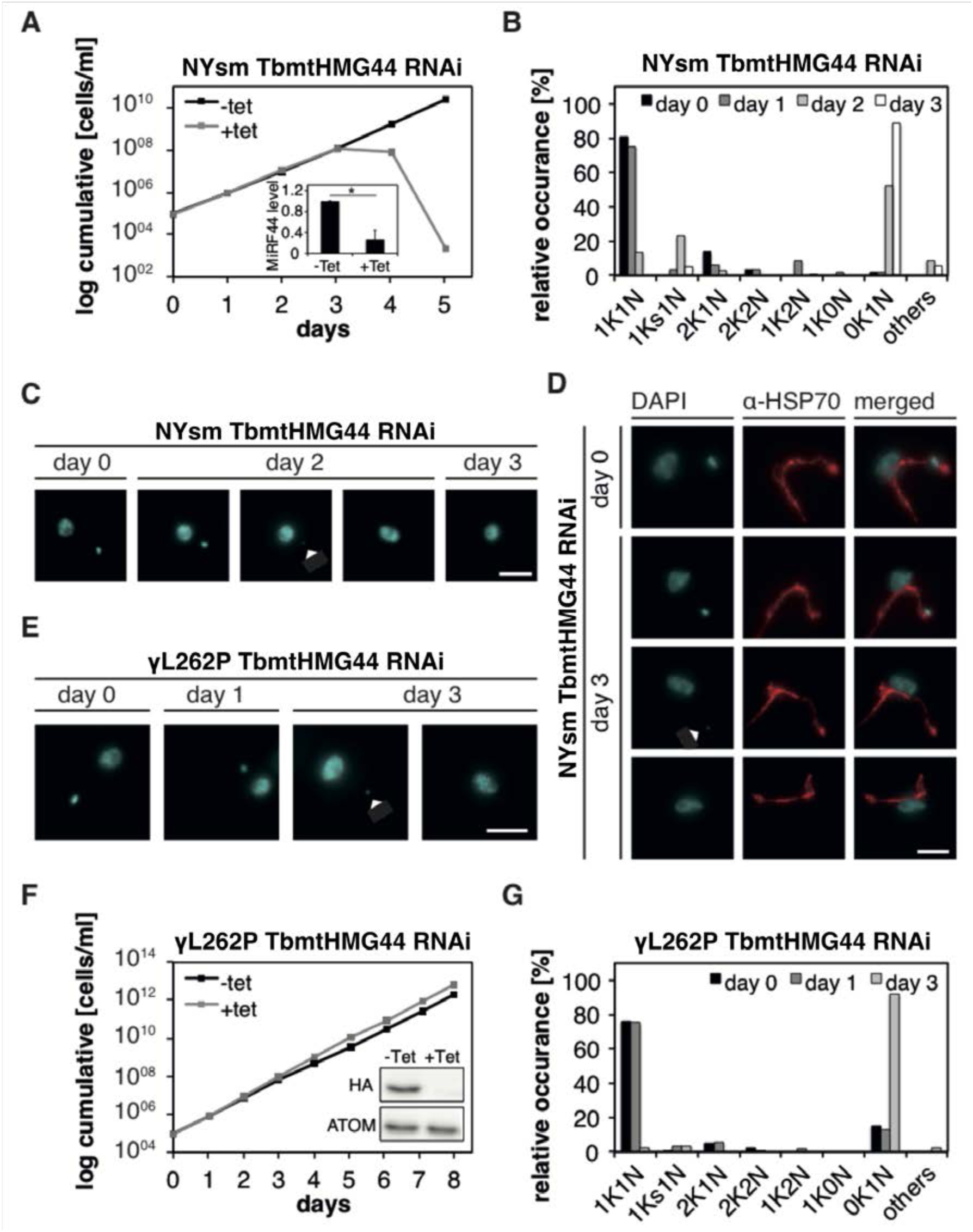
Phenotype upon TbmtHMG44 depletion. **A)** Growth curve of TbmtHMG44 uninduced (-tet) and induced (+tet) NYsm cells. Inset: qPCR in triplicates for uninduced and 3 days induced cells (N=3) Normalized to α-tubulin. Uninduced cells equal one. The p-value was calculated to perform significance measurements (two-tailed heteroscedastic t-test; * 0.01<p≤0.05). **B)** Cell cycle stages at different time points after TbmtHMG44 depletion (N≥104). **C)** Representative images of uninduced (day 0) and induced (day 2, day 3) cells. Arrowhead points to a small kDNA. Cyan: DAPI (DNA). **D)** Mitochondria (red, anti-mtHSP70) in uninduced and 3 days induced cells. Cyan: DAPI (DNA). **E)** Representative images of uninduced and induced cells (day 1, day 3). Arrowhead points to a small kDNA. Cyan: DAPI (DNA). **F)** Growth curve in the γL262P cell line upon depletion. Inset: western blot shows the protein abundance of TbmtHMG44 in uninduced and 3 days induced cells. ATOM40 was used as a loading control. **G)** Cell cycle stages at 0, 1 and 3 days after TbmtHMG44 depletion (N≥110). Scale bar 3 µm.

**Figure S9:**
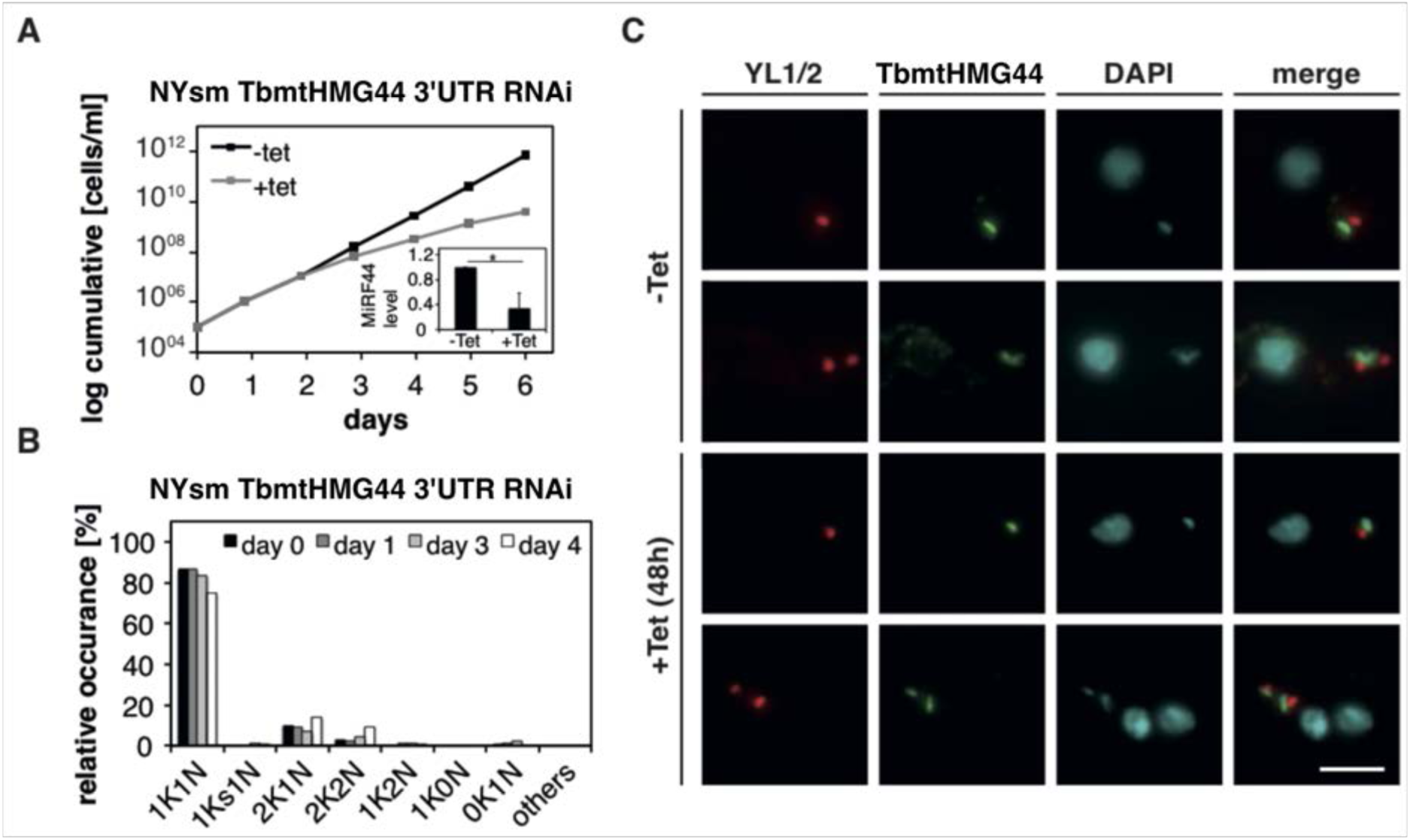
Functionality study of the HA-tagged TbmtHMG44 in BSF cells. **A)** Growth curve of RNAi against the 3’UTR of TbmtHMG44 uninduced (-tet) or induced (+tet) TbmtHMG44 HA-tagged cells. Inset: qPCR in triplicates for uninduced and three days induced cells (N=3). Normalized to tubulin. Uninduced cells equal one. The p-value was calculated to perform significance measurements (two-tailed heteroscedastic t-test; * 0.01<p≤0.05). **B)** Cell cycle stage at 0, 1, 3 and 4 days after RNAi induction (N≥86). **C)** Representative images of uninduced and 2 days induced cells. Red: YL1/2 (basal body), green: HA (TbmtHMG44), cyan: DAPI (DNA). Scale bar 3 µm.

**Figure S10:**
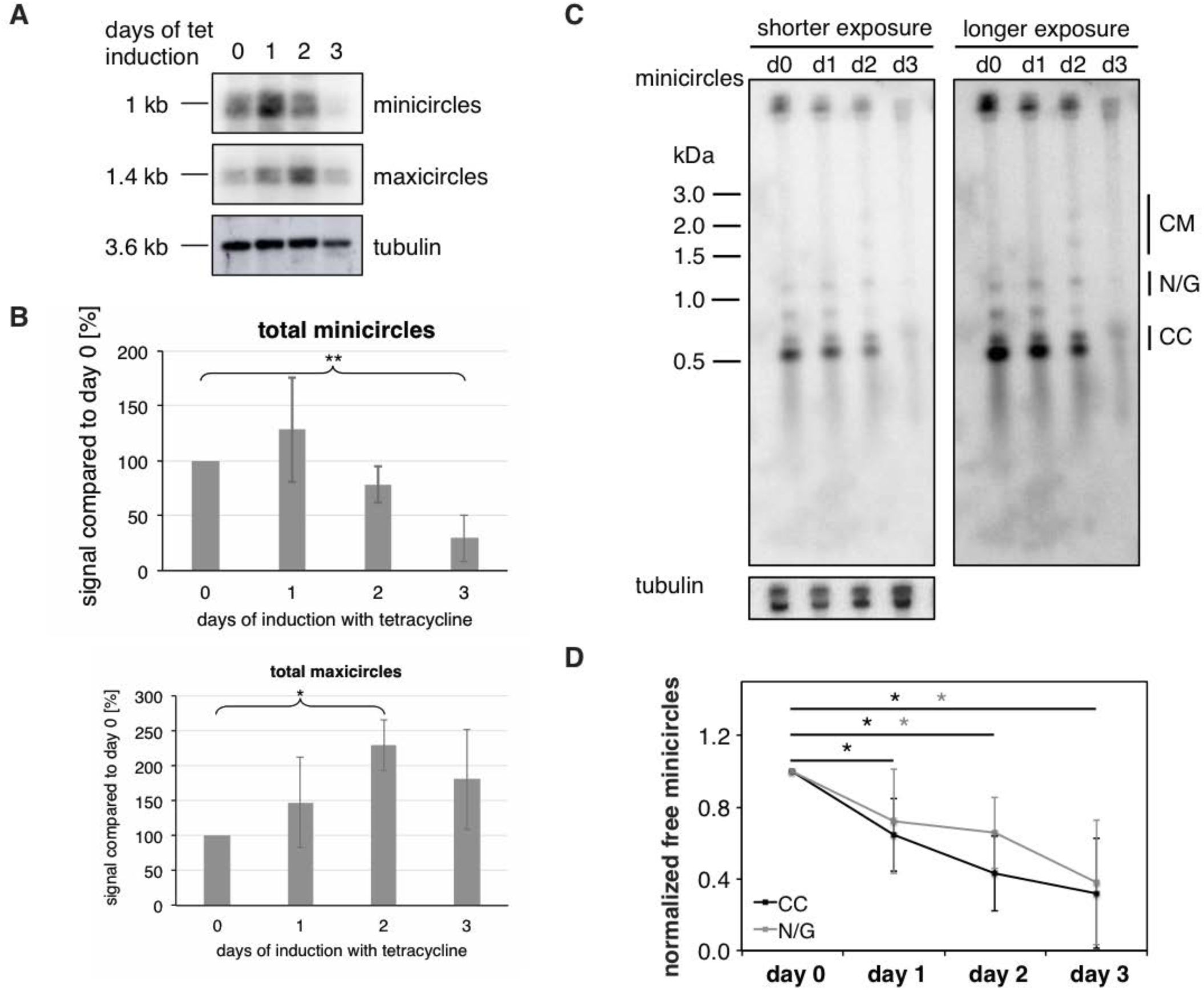
Southern blot analysis of TbmtHMG44 RNAi. **A)** Southern blot probing for mini and maxicircles after TbmtHMG44 depletion. Digested DNA of TbmtHMG44 RNAi cells, after induction of TbmtHMG44 RNAi using tetracycline, is loaded on a 1% agarose gel to detect total minicircles and maxicircles by Southern blot. Tubulin was used as a loading control. **B)** Quantification of the total minicircles (n = 6) and maxicircles (n = 5) normalized to tubulin (n = 5). **C)** Effect on free minicircle replication intermediates. Undigested DNA was purified and loaded on a 1.5 % agarose gel containing ethidium bromide. After transferring onto a PVDF membrane, kDNA was detected by using a minicircle probe. Tubulin was used as a loading control. CC: covalently closed; N/G: nicked/gapped; CM: catenated minicircles. **D)** Quantification of CC and N/G minicircles normalized to tubulin and day 0 equals one (N = 4). The p-values were calculated to perform significance measurements (two-tailed heteroscedastic t-test; * 0.01 < p ≤ 0.05; ** 0.001 < p ≤ 0.01; *** p ≤ 0.001).

**Figure S11.**
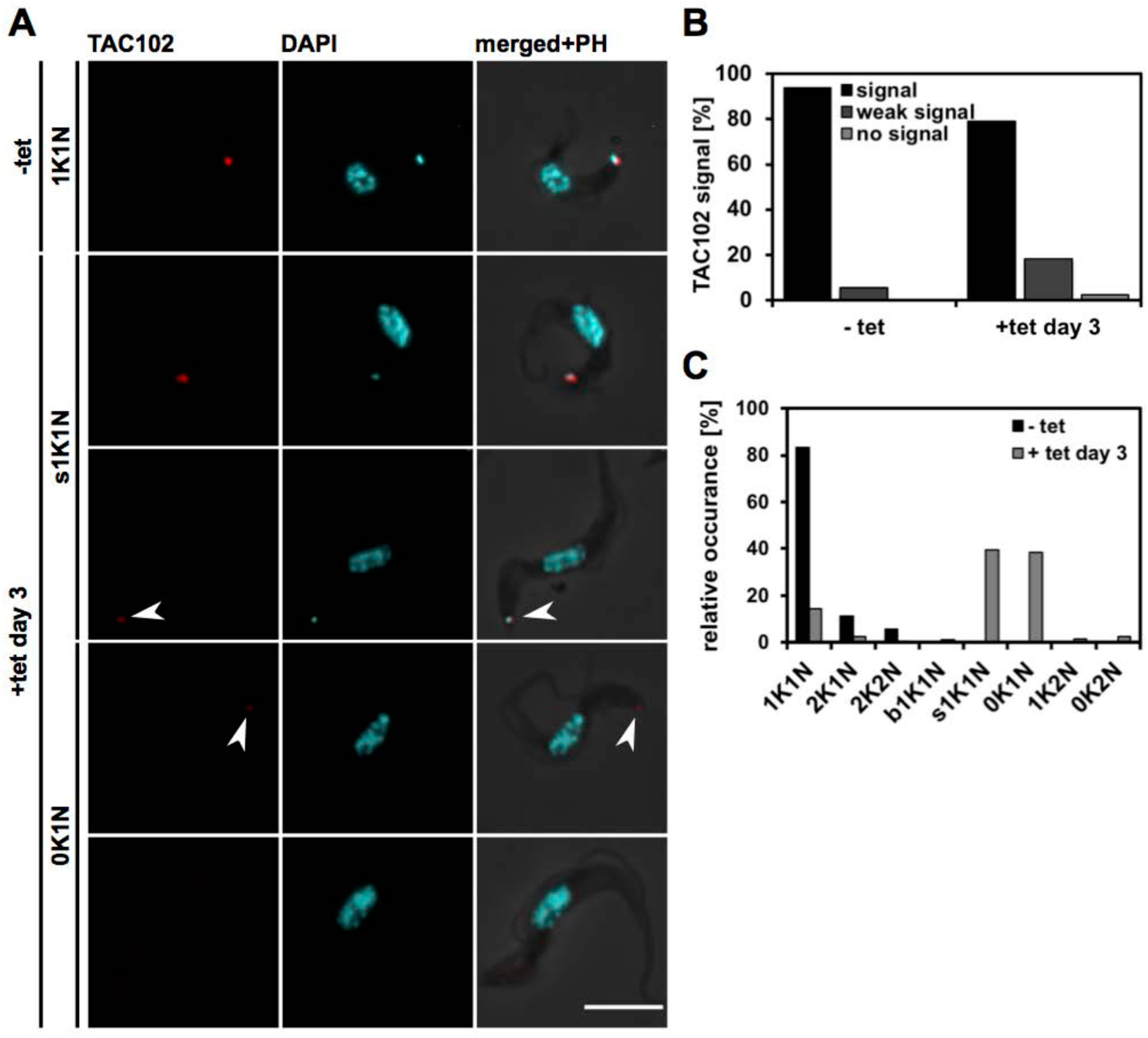
Effect of Tb927.11.16120 depletion on TAC102 in BSF cells. **A)** The same cell line as described in the previous figure was used. Immunofluorescence microscopy images were generated as described in the previous figures. TAC102 and the kDNA were detected with the same reagents as described in the previous figures. **B)** Quantification of TAC102 signal in uninduced cells (-tet) and cells at day three post induction (+tet days 3) (n≥228 for each condition). **C)** Quantification of the relative occurrence of kDNA discs and nuclei (n≥196 for each condition). bK, big kDNA; K, kDNA; N, nucleus; PH, phase contrast; sK; small kDNA. Scale bar: 5 μm.

**Figure S12:**
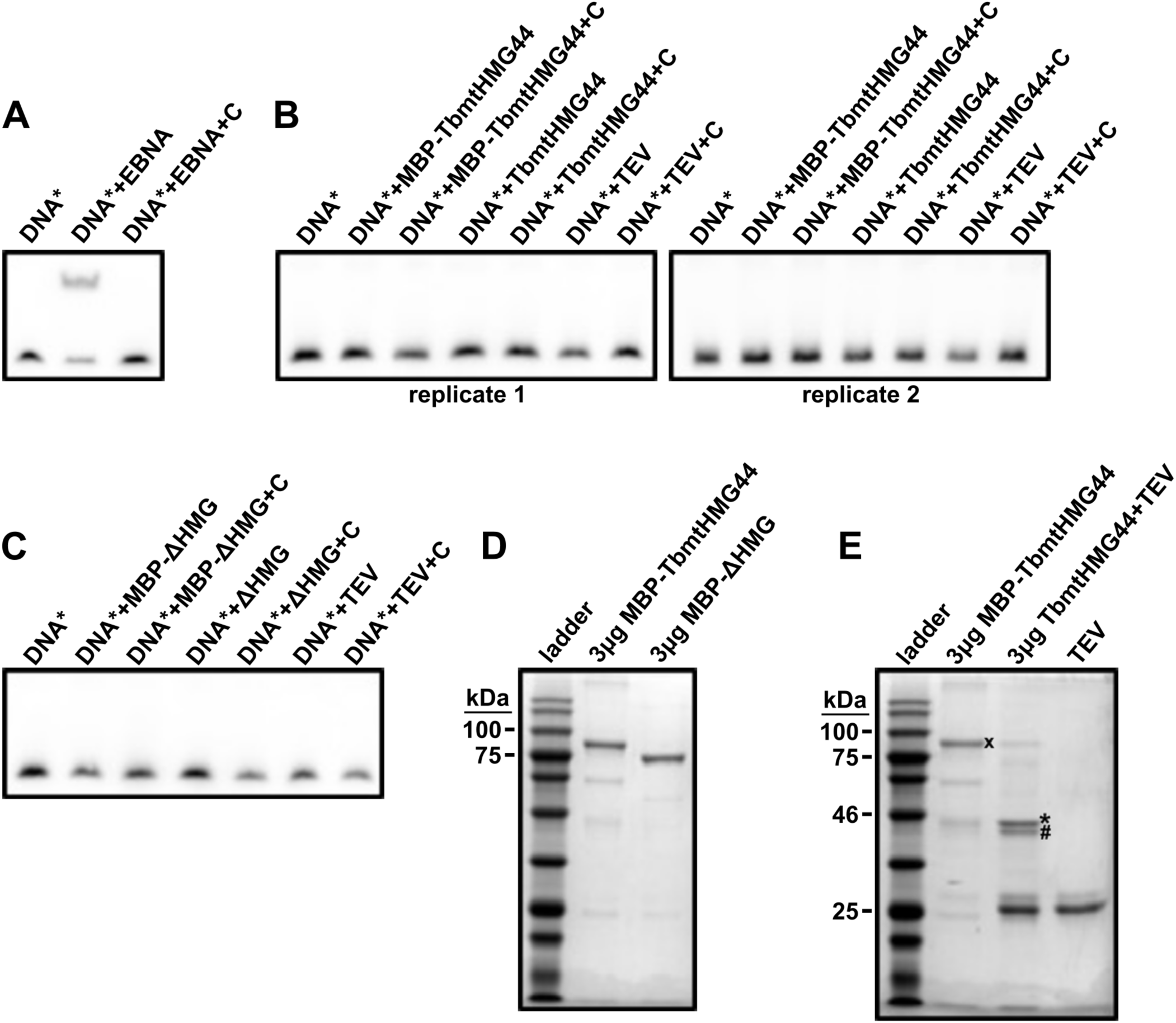
Analysis of putative TbmtHMG44 nucleic acid binding activity using electro mobility shift assays (EMSA) with recombinant MBP-TbmtHMG44. **A)** EMSA performed with EBNA DNA as bait and EBNA nuclear protein extract as positive control for the test of the kit performance. **B)** EMSA with recombinant MBP-TbmtHMG44 containing an N-terminal MBP tag and recombinant TbmtHMG44 where the tag was cleaved away by TEV (TbmtHMG44). As negative control we performed an EMSA with TEV. **C)** EMSA with mutated recombinant MBP-TbmtHMG44 missing the HMG box (MBP-ΔHMG) and same mutated version where the MBP was cleaved off as described above (ΔHMG). TEV control was performed as described above. For the experiments in B) and C) we used a region of the conserved minicircle sequence as bait. **D)** Coomassie stained gel showing purified recombinant MBP-TbmtHMG44 and recombinant TbmtHMG44 missing the HMG box (MBP-ΔHMG). **E)** Coomassie stained gel showing the TEV cleavage exemplified by recombinant MBP-TbmtHMG44. TEV,; tobacco etch virus protease; x, MBP-5020; *, 5020; #, MBP

## Supplementary Tables

**Table S1:**
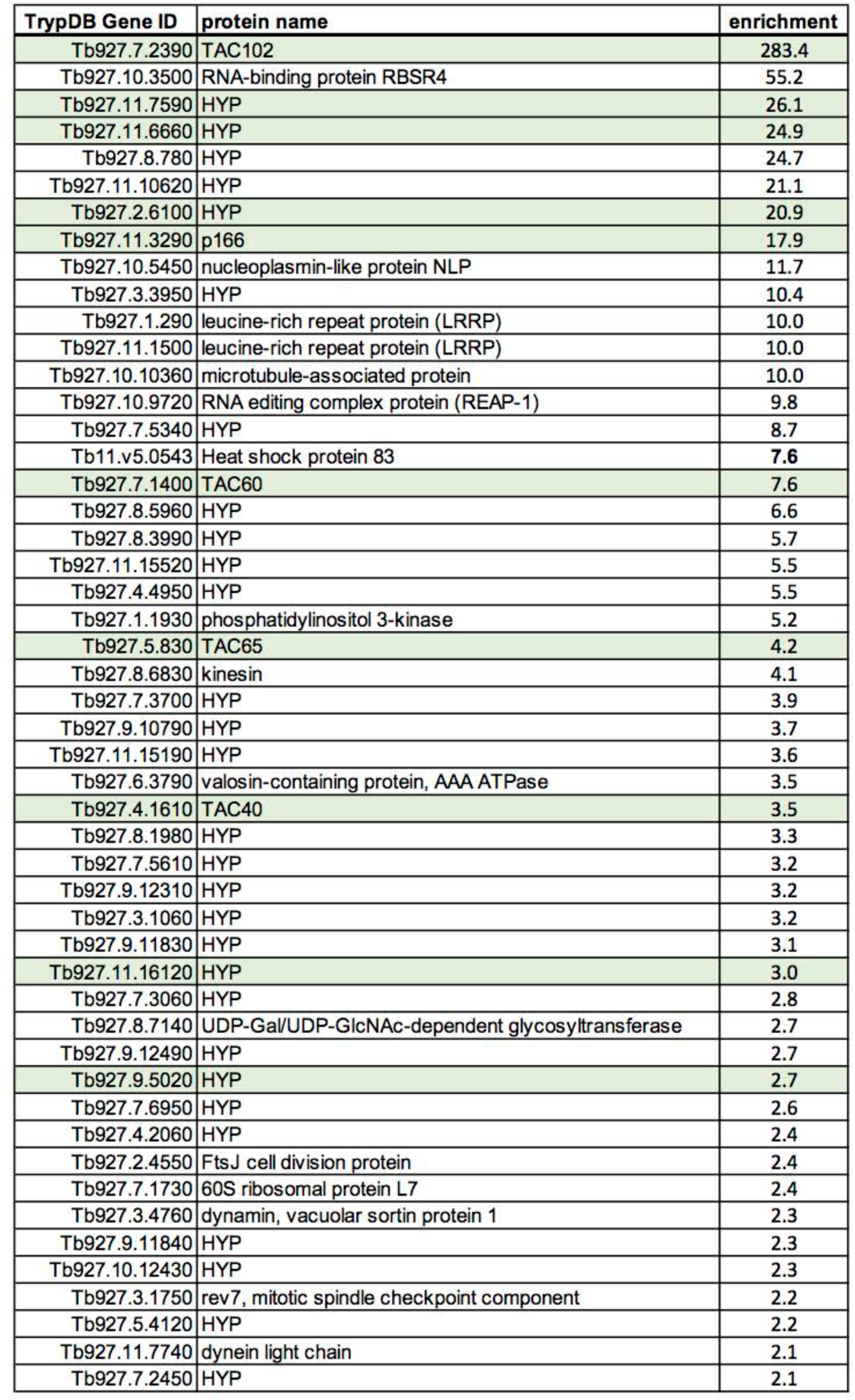
Candidates of the YFP-TAC102 IP. Top 50 proteins enriched in the YFP-TAC102 IP. The enrichment of the proteins was calculated by the spectral index ratio of the eluate to the flow-through. HYP = hypothetical protein, conserved.

**Table S2:**
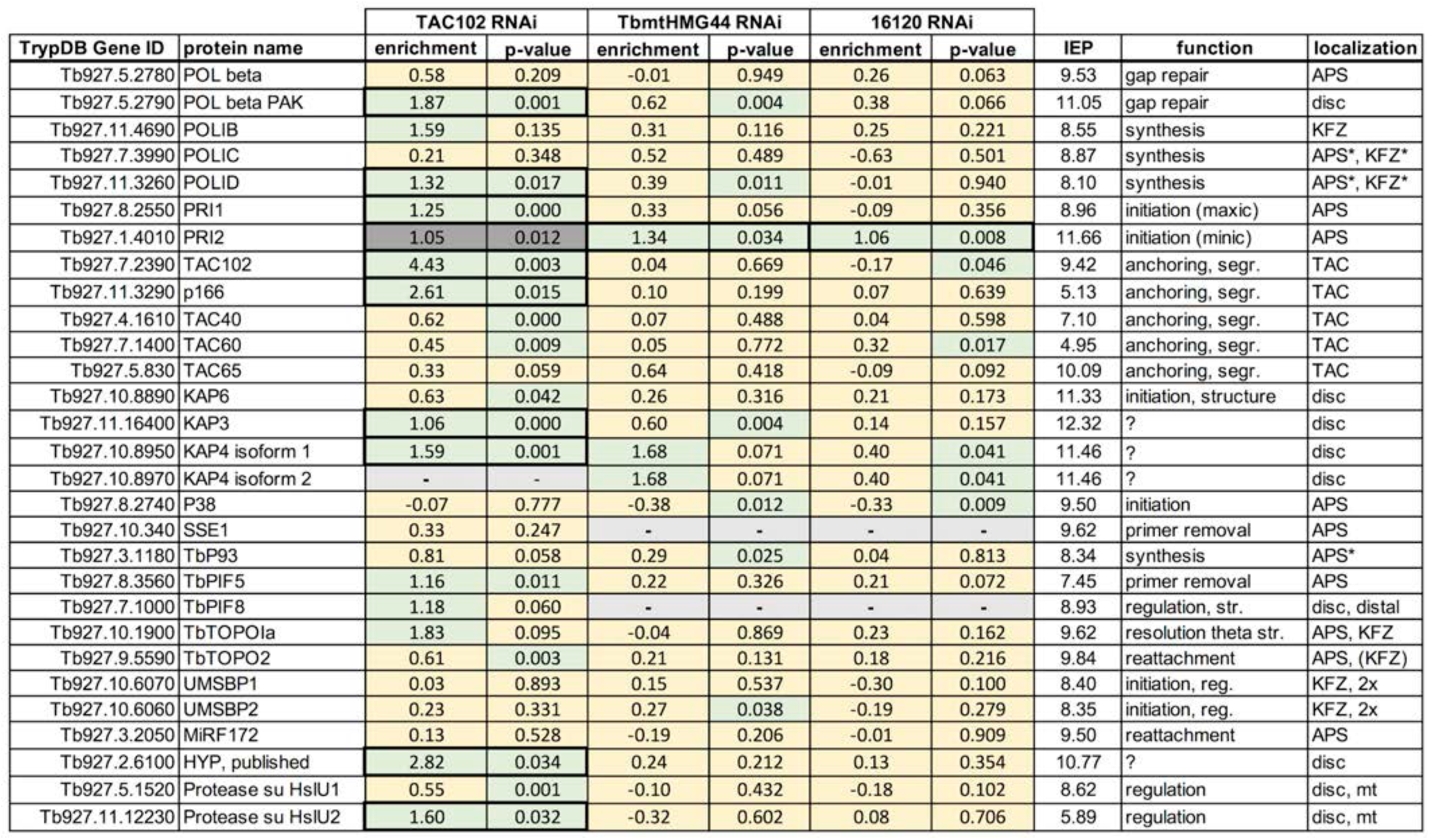
kDNA replication/segregation factors identified in the flagellar extract mass spectrometry analysis. Yellow unchanged abundance (log_2_), green changed abundance (thresholds: p-value <0.05, fold change −1< log_2_ >1). Grey values were calculated based on less than four replicates. APS, antipodal sites; disc, localization within kDNA network; distal, localization at the basal body distal phase of the network; KFZ, kinetoflagellar zone; maxic, maxicircle; minic, minicircle; mt, mitochondrion; reg, regulation; segr., segregation; *, localization only observed during kinetoplast S phase

